# A Novel ψ-χ Fusion Protein for Unravelling the Contributions of χ to DNA Replication and Repair

**DOI:** 10.1101/2025.10.15.682636

**Authors:** Kaylie A. Padgett-Pagliai, Elise Wimer, Jacob D. Grant, Matthew J. Petrides, Elijah S. P. Newcomb, Vincent A. Sutera, Susan T. Lovett, Linda B. Bloom

## Abstract

Faithful DNA replication in *Escherichia coli* requires the DNA polymerase III holoenzyme (DNA pol III HE) and its clamp loader complex, which couples processive DNA synthesis with β-clamp loading. The clamp loader accessory subunits χ and ψ link single-stranded DNA-binding protein (SSB) to the replisome, stabilizing replication on SSB-coated templates through interactions with the SSB C-terminal tail. Chi has also been implicated in tolerance to the chain-terminating nucleoside analog azidothymidine (AZT), though whether this function depends on χ within DNA pol III HE or on its independent interaction with the YoaA helicase remains unclear. To address this, we engineered ψ-χ fusion proteins with flexible glycine-serine linkers to tether the two subunits while preserving folding and activity. Both fusions were biochemically competent, supporting ATP hydrolysis and clamp loading on SSB-coated DNA. In vivo, however, neither the ψ-GS12-χ fusion nor expression of a ψχ operon restored AZT tolerance in Δ*holC* cells, whereas expression of χ alone was sufficient. Fusion expression impaired growth in both wild-type (WT) and Δ*holC* backgrounds, a phenotype alleviated by disrupting χ-SSB binding. These findings support a model in which χ must dynamically engage SSB and YoaA outside of the clamp loader to promote AZT tolerance, highlighting the importance of regulated χ-SSB interactions in genome maintenance.

## INTRODUCTION

The fidelity of DNA synthesis and repair reactions is critical for all cells to overcome replication errors. In *Escherichia coli*, most DNA synthesis is accomplished by DNA polymerase III holoenzyme (DNA pol III HE). This highly processive enzyme coordinates replication of both the leading and lagging DNA strands at the replication fork.

The DNA pol III HE (reviewed in (1, 2) consists of two distinct components: the pol III core polymerase and the clamp loader complex (Figure 1). The pol III core polymerases (αεθ) actively synthesize the leading and lagging strands. The clamp loader complex (τ_2_[τ/γ]δδ’ψχ) coordinates the replisome by loading the processive β-sliding clamp onto DNA. Active clamp loaders contain three copies of the DnaX subunit, existing in two forms within the cell: the full-length form (τ), and the truncated form (γ), generated by a translational frameshift (3–5). The τ subunit contains an extended C-terminal domain that binds the DNA polymerase core and the DnaB helicase, physically coupling polymerase and helicase activities at the replication fork (3, 6). In contrast, γ lacks this domain and cannot tether the polymerase or DnaB, resulting in clamp loader with different structural and functional properties.

**Figure 1:**
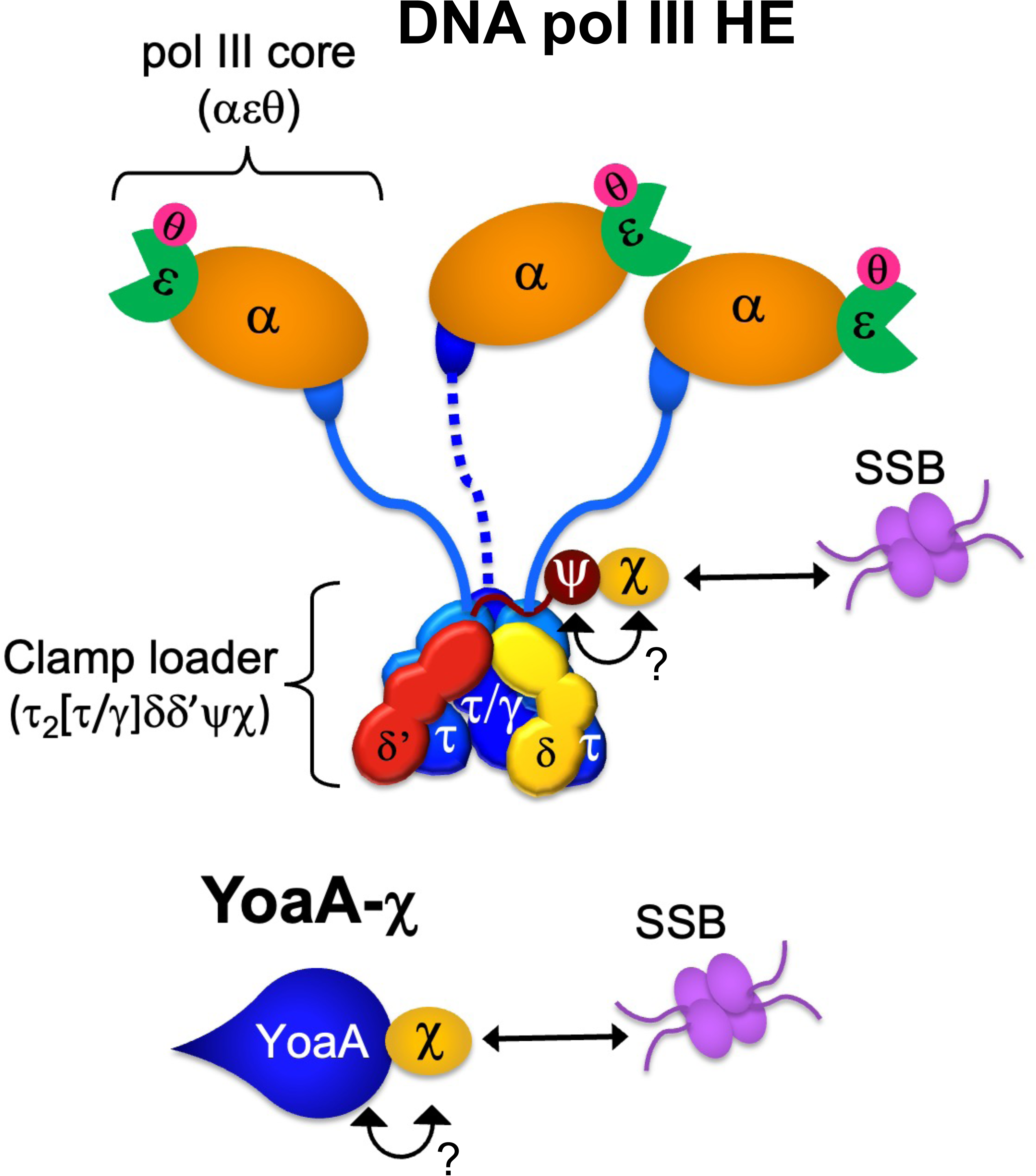
Schematic diagram of the DNA pol III HE (without the β-clamp) and YoaA-χ. Interactions between χ and SSB are needed for AZT tolerance as indicated by arrows, but which χ complex plays an important role is unknown.

Active clamp loaders can be made using any combination of DnaX subunits (7). The γ complex clamp loader used in this study (γ_3_δδ’ψχ) lacks the C-terminal region of τ, which binds the core polymerase but is subject to proteolysis during purification in the absence of the polymerase. Multiple forms of the γ complex clamp loader can exist in vivo depending on the relative expression of τ and γ. The ability to assembly both types of clamp loaders enables dynamic regulation of replisome composition in response to cellular needs (3, 6, 8).

Within the clamp loader, the ψχ complex functions as an accessory module to the DNA pol III HE and can exist as a heterodimer. Although χ is not essential for polymerase or clamp loader activity, it is the only known subunit of DNA pol III HE that makes direct physical and functional contact with the single-strand DNA-binding protein (SSB). Psi binds the clamp loader and χ, thereby linking SSB to DNA pol III HE and enhancing replication efficiency on SSB-coated DNA (Figure 1) (9–11). Chi is also important for efficiently loading clamps on DNA bound by SSB, emphasizing that SSB-clamp loader interactions hold intrinsic physiological significance even in the absence of the full DNA pol III HE (12).

In addition to SSB, χ has been shown to interact with YoaA, an XPD/Rad3-like helicase (Figure 1) (13, 14). The YoaA-χ complex is described as the likely physiological form of the helicase. Although *holC* (encoding χ) is not essential, *holC*-null mutants exhibit poor growth and elevated rates of genetic rearrangements (15). Duplication of *ssb* is known to suppress the growth defects of both *holC-* and *holD*-null mutants, which otherwise rapidly acquire secondary genetic suppressor mutations (16). Interestingly, deletion of *yoaA* in a *holC*-mutant background improves growth and viability, suggesting that YoaA is toxic in the absence of χ (17). Both YoaA and χ contribute to the maintenance of genome integrity, including roles in repairing methyl methanesulfonate (MMS)-induced lesions (18). Moreover, *yoaA* and *holC*, along with SSB-χ interactions, confer tolerance to 3′-azido-3′deoxythymidine (AZT), a thymidine analog that blocks DNA synthesis by chain termination (17, 19).

Consistent with this, mutations in YoaA that disrupt interactions with χ compromise AZT tolerance (13). Notably, the interaction between χ and ψ, which provides the only known connection to DNA pol III, is mutually exclusive with the interaction between χ and YoaA (13). This raises the question of whether the role of χ in AZT tolerance depends on its association with DNA pol III, YoaA, or both (Figure 1).

The overarching goal of this study is to develop a tool that can be used to resolve the distinct contributions of χ to DNA replication and DNA damage tolerance. To do this, we engineered ψ-χ fusion proteins that can be uniquely incorporated into DNA pol III HE, leaving other pools of χ available for non-polymerase functions. We designed two fusions containing flexible glycine-serine linkers, ψ-GS12-χ and ψ-GS8-χ, and biochemically characterized them as unique complexes and within the context of the γ complex clamp loader. The ψ-GS12-χ fusion was further tested in vivo to determine whether it could rescue the growth defect of a χ deletion strain and whether it contributes to AZT tolerance. This work establishes a stable and functional, novel ψ-χ fusion protein that operates within both the clamp loader and the DNA pol III HE. Importantly, the ψ-χ fusion provides a tool to distinguish between polymerase-associated χ from helicase-associated χ, enabling future studies to examine the dual roles of χ in DNA replication and repair.

## MATERIAL AND METHODS

### Protein Models

Structural models for Psi-GS12-Chi and Psi-GS8-Chi were made using AlphaFold Colab (https://colab.research.google.com/github/deepmind/alphafold/blob/main/notebooks/AlphaFold.ipynb) on January 20^th^, 2023, and January 31^st^, 2023, respectively. AlphaFold Colab runs a simplified version of AlphaFold v2.3.1 (20).

### Strains and Media

The strains used in this study are listed in Table 1. *E*. *coli* strains were cultured in either lysogeny broth (LB) or Terrific Broth (TB) as specified. Culture media were amended with the appropriate concentration of antibiotics as recommended for each genetic vector (Table 1).

**Table 1.**
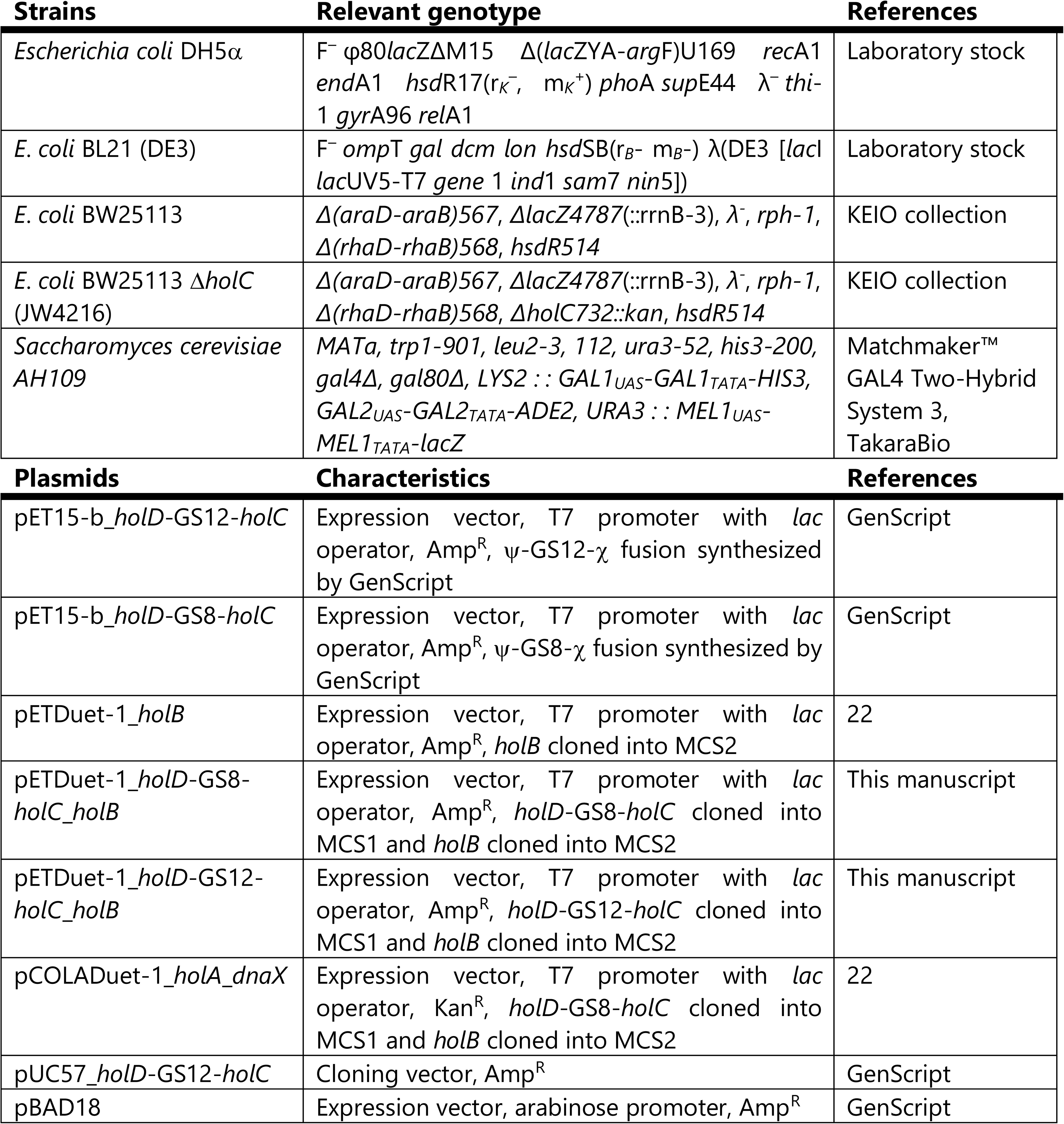
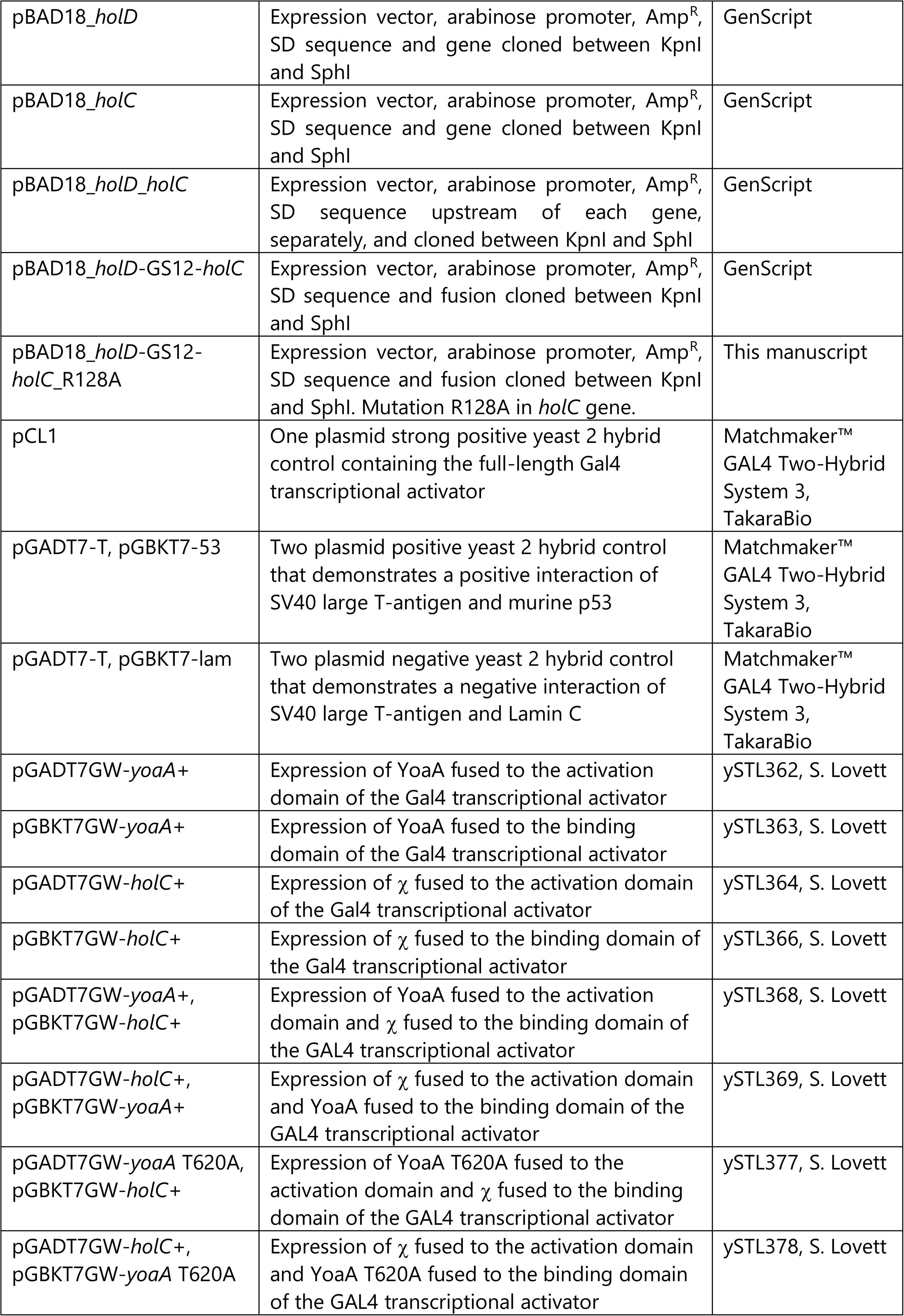
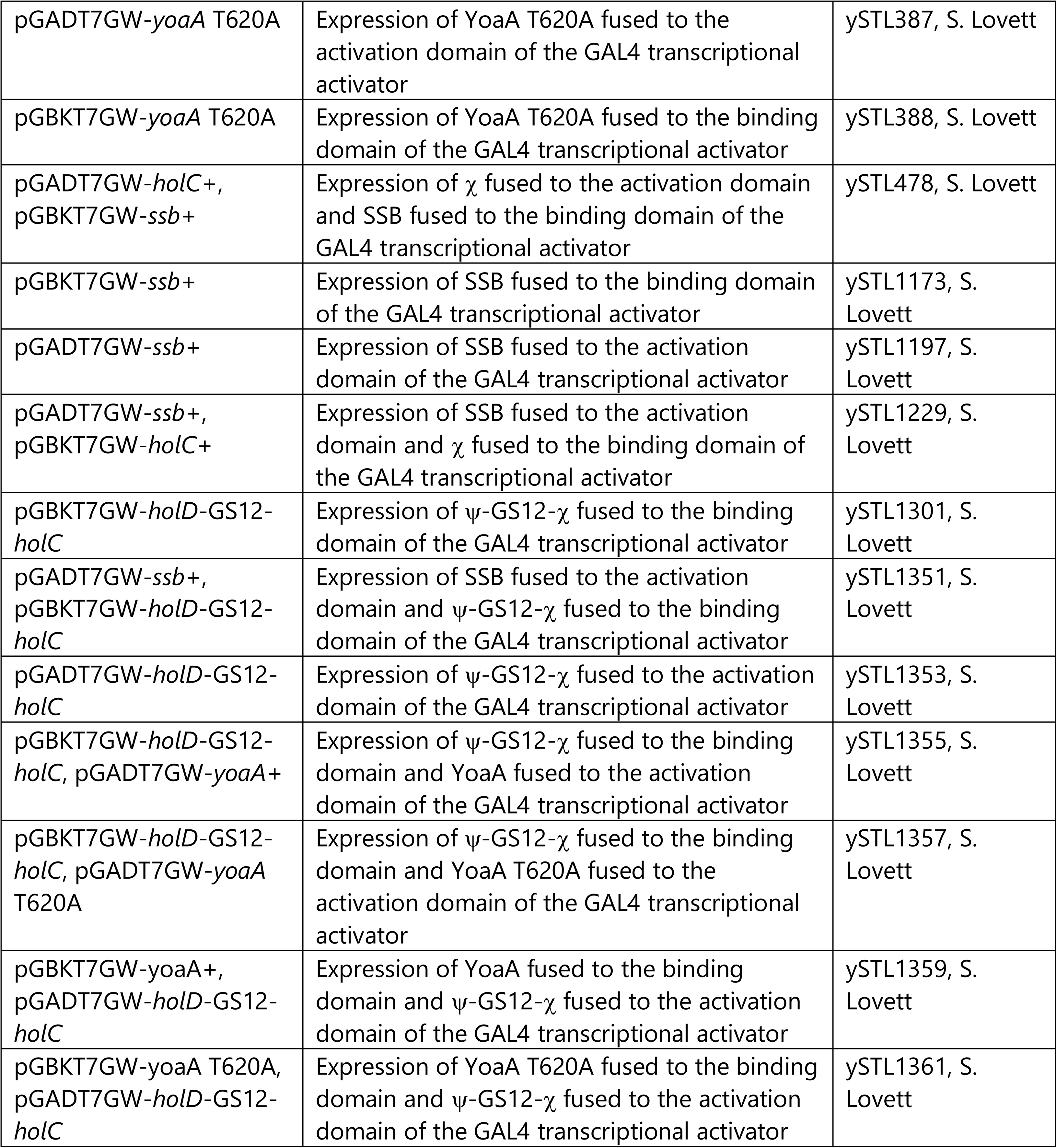
Strains and plasmids used in this study.

### Cloning

The amino acid and nucleotide sequences for the GS8 and GS12 fusion are listed in Table 2.

**Table 2.**
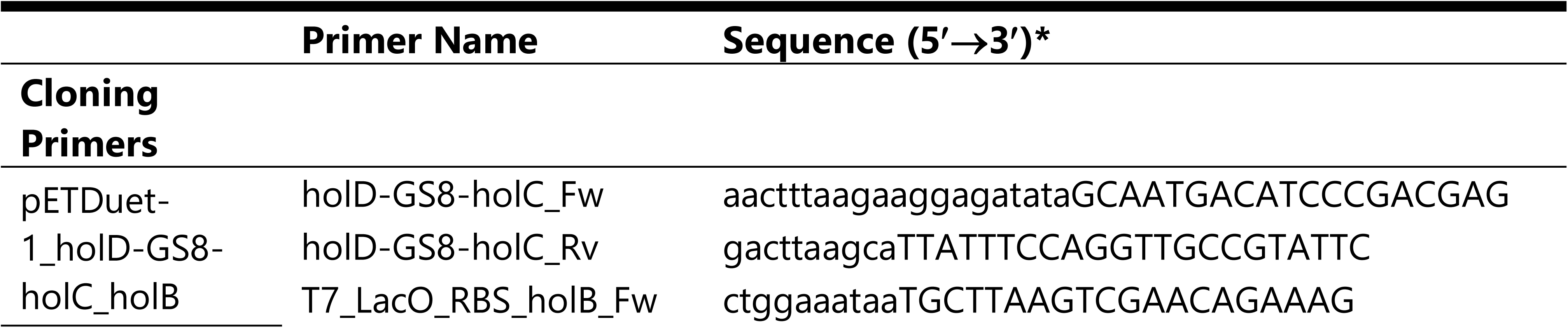
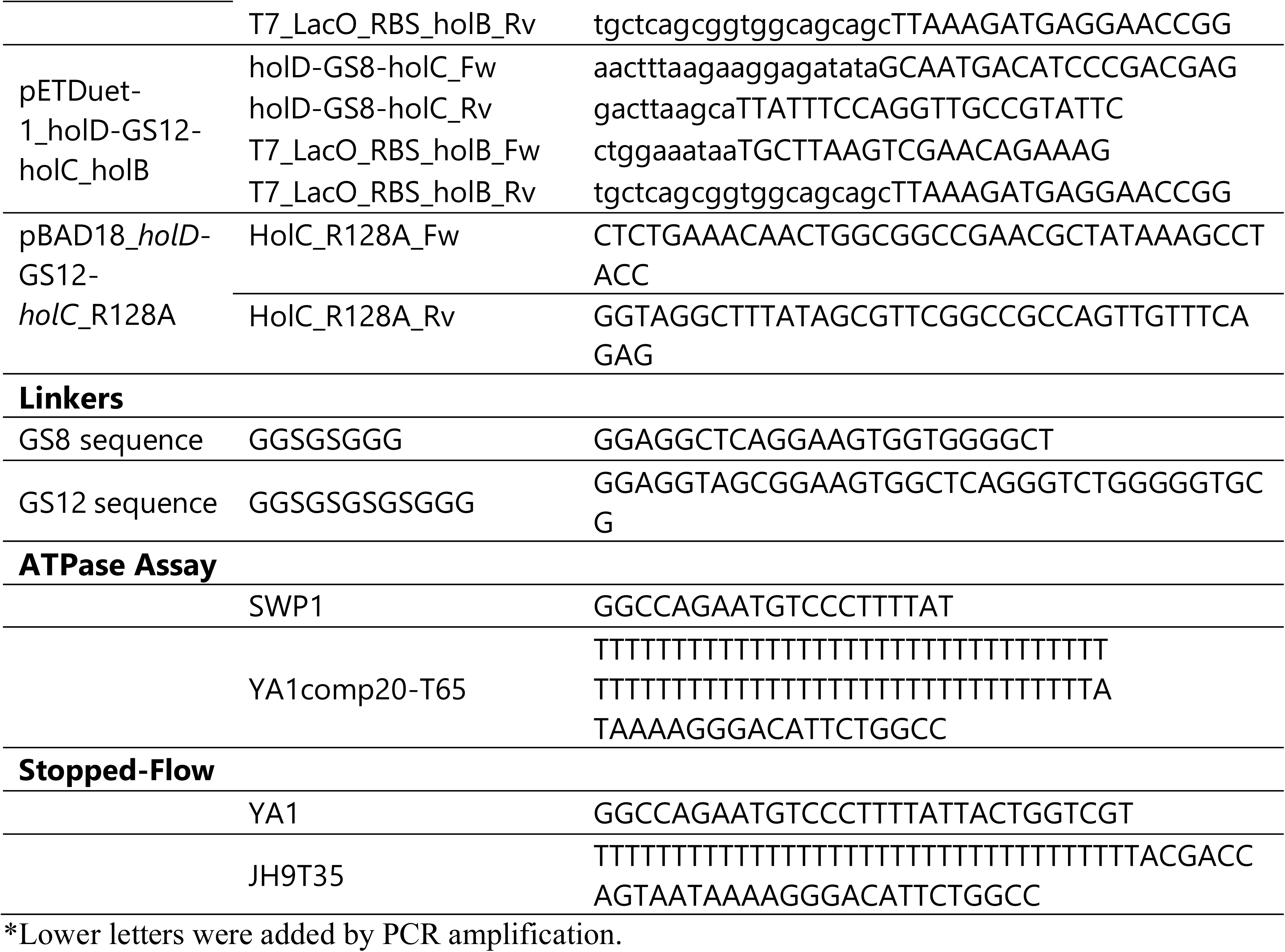
Oligonucleotides by use.

The *holD*-GS12-*holC* was originally synthesized and cloned by GenScript into pUC57 between XbaI/BamHI. The terminal stop codon of *holD* was omitted, and the nucleotide sequence for GS12 was inserted immediately upstream of *holC*. pUC57_*holD*-GS12-*holC* was then transformed into *E*. *coli* DH5α and the plasmid extracted (Quiagen). To express *holD*-GS12-*holC* with *holB*, the plasmid was digested using XbaI (New England Biolabs, NEB) and BamHI (NEB). XbaI-*holD*-GS12-*holC*-BamHI was then ligated into MCS1 of pETDuet-1_*holB* using the sites XbaI/BamHI, creating pETDuet-1_*holD*-GS12-*holC*_*holB*.

GenScript also synthesized *holD*-GS8-*holC* and *holD*-GS12-*holC* in pET-15b using the restriction sites NcoI and BamHI to express the fusion proteins with a 6X N-terminal histidine tag. The terminal stop codon of *holD* was omitted, and the nucleotide sequence for GS8 or GS12 was inserted immediately upstream of *holC*.

To clone *holD*-GS8-*holC* into pETDuet-1, Gibson Assembly^®^ (NEB) was designed according to the manufacturer’s instructions. Fragments for *holD*-GS8-*holC* and *holB* were PCR amplified using Q5 High Fidelity Master Mix (NEB) following the manufacturer’s instructions. *holD*-GS8-*holC* was amplified from pET15-b_*holD*-GS8-*holC* vector using the primers *holD*-GS8-*holC*_Fw and *holD*-GS8-*holC*_Rv listed in Table 2. *holB* was amplified from pETDuet-1_*holB* using the primers T7_LacO_RBS_*holB*_Fw and T7_LacO_RBS_*holB*_Rv listed in Table 2. pETDuet-1 was cut with AvrII (NEB) and NcoI (NEB) and gel purified (QIAquick^®^ Gel Extraction Kit, QIAGEN). The Gibson Assembly^®^ Cloning Kit (NEB) was utilized according to the manufacturer’s instructions to assemble fusion proteins into MCS1 of pETDuet-1 and *holB* into MCS2, creating pETDuet-1_*holD*-GS8-*holC*_*holB*.

Complementation vectors were generated using Genscript. The Shine Dalgarno (SD) sequence and spacer (AGGAGGaattc), based upon previous literature, were inserted upstream of each gene in the pBAD18 vector, and the complete constructs were cloned between the *KpnI* and *SphI* sites (21). The following inserts were cloned: *holD*, *holC*, a *holD*_*holC*_operon, with an SD sequence and spacer preceding each gene, and holD-GS12-holC. The plasmid pBAD18_*holD*-GS12-*holC* was used as the template to generate the R128A mutant by site-directed mutagenesis using the primers HolC_R128A_Fw and HolC_R128A_Rv listed in Table 2. Mutagenesis was performed using the Q5^®^ Site-Directed Mutagenesis Kit (NEB) according to the manufacturer’s instructions.

### Protein Expression and Purification

The γ complex clamp loader (γ_3_δδ’ψχ subunits) and β-clamp were expressed and purified as described previously (12, 22–24). Unlabeled protein concentrations were determined by measuring the absorbance at 280 nm under native conditions using extinction coefficients of 220,050 M^-1^ cm^-1^ for γ complex (25) and 14,700 M^-1^ cm^-1^ for β (23). All other protein concentrations were measured using a Bradford assay at an absorbance of 595 nm.

#### Expression of ψ-GS8-χ and ψ-GS12-χ. For ψ-GS8-χ, E. coli

BL21 (DE3) cells were transformed with pET-15b_*holD*-GS8-*holC*. Cells were suspended in Terrific Broth supplemented with Carbenicillin 100 ng ml^-1^ (Carb100), grown at 37°C, shaking at 250 RPM until OD_600_ = 0.6. IPTG was added at a final concentration of 1 mM to induce protein expression, and cells were incubated at 37°C, shaking at 250 RPM, for 2 h. Cells were subsequently pelleted through centrifugation at 4°C, 6800 RCF for 30 min, decanted, and stored at −80°C. For ψ-GS12-χ, *E*. *coli* BL21 (DE3) cells were transformed with pET-15b_*holD*-GS12-*holC*. Cell growth and protein induction were done as previously described for ψ-GS8-χ.

#### Purification of ψ-GS8-χ and ψ-GS12-χ

Pellets of ψ-GS8-χ were resuspended in cell lysis/low imidazole buffer (50 mM Tris-HCl pH 7.5, 500 mM NaCl, 75 mM imidazole, 10% glycerol, 2 mM DTT, Pierce Protease Inhibitor Tablets – EDTA Free (ThermoScientific, Waltham, MA, USA)) and lysed by two passes through a French Press cell disruptor. Lysate was clarified by centrifugation at 12,000 RCF for 45 min, and supernatant was applied to a HisTrap FF column (Cytiva, Marlborough, Massachusetts, USA). After washing with low imidazole buffer, bound protein was eluted using a linear imidazole gradient (75 mM to 500 mM). ψ-GS8-χ eluted in two distinct peaks, at approximately 210 and 310 mM imidazole. Elution fractions were pooled and dialyzed overnight against dialysis buffer (50 mM Tris-HCl pH 7.5, 500 mM NaCl, 20% glycerol, and 2 mM DTT), followed by a second dialysis in SP HP low salt buffer (50 mM Tris-HCl pH 7.5, 20 mM NaCl, 20% glycerol, and 2 mM DTT). The first elution peak retained from the nickel column was then applied to an SP HP column, where precipitation occurred during binding. Protein was recovered by elution with SP HP high salt buffer (50 mM Tris-HCl pH 7.5, 500 mM NaCl, 20% glycerol, and 2 mM DTT), re-purified through nickel affinity chromatography, and dialyzed sequentially in dialysis buffer and then overnight into storage buffer (50 mM Tris-HCl pH 7.5, 50 mM NaCl, 20% glycerol, and 2 mM DTT). The second peak from the nickel column did not go through the SP HP column and was dialyzed directly into storage buffer.

Pellets of ψ-GS12-χ were resuspended in cell lysis/low imidazole buffer and lysed as described above. Lysate was clarified at 12,000 RCF for 45 min, and supernatant loaded onto a HisTrap FF column (Cytiva). After washing with low imidazole buffer, protein was eluted using a linear imidazole gradient (40-500mM), eluting at approximately 210 mM imidazole. Fractions were dialyzed overnight in dialysis buffer and SP HP low salt buffer as described above. Protein was eluted using a linear salt gradient (20mM to 200mM), with ψ-GS12-χ eluting at two peaks, approximately 135 and 180 mM NaCl. Peak fractions were pooled separately, dialyzed overnight into storage buffer, and stored at −80°C as described above.

#### Expression of Clamp Loaders with ψ-GS8-χ and ψ-GS12-χ

For clamp loader containing ψ-GS8-χ, *E*. *coli* BL21 (DE3) cells were co-transformed with pETDuet-1_*holD*-GS8-*holC*_*holB* for the expression of ψ-GS8-χ and δ′, as well as pCOLADuet-1_*holA*_*dnaX* (24) for the expression of δ and γ. Cells were suspended in Terrific Broth supplemented with Carb100 and Kanamycin 50 ng ml^-1^ and grown at 37°C, shaking at 250 RPM until OD_600_ = 0.6. IPTG was added at a final concentration of 1 mM to induce protein expression and incubated at 37°C, shaking at 250 RPM, for 3 h. Cells were subsequently pelleted through centrifugation at 4°C, 6800 RCF for 30 min, decanted, and stored at −80°C. For the clamp loader containing ψ-GS12-χ, *E*. *coli* BL21 (DE3) cells were co-transformed with pETDuet-1_*holD*-GS12-*holC*_*holB* for the expression of ψ-GS12-χ and δ′, as well as pCOLADuet-1_*holA*_*dnaX* for the expression of δ and γ. Cells were grown and induced as stated for clamp loader containing ψ-GS8-χ.

#### Purification of Clamp Loaders with ψ-GS8-χ and ψ-GS12-χ

Pellets of clamp loader ψ-GS8-χ were resuspended in cell lysis/low salt buffer (50mM Tris-HCl pH 7.5, 100mM NaCl, 5% glycerol, 2mM DTT, Pierce Protease Inhibitor Tablets – EDTA Free (ThermoScientific) and lysed as described above. Lysate was clarified at 12,000 RCF for 45 min, and the supernatant loaded onto an SP HP column (Cytiva). Because the clamp loader does not bind SP HP at 100mM NaCl, the flow-through was dialyzed overnight against Q HP low salt buffer (20 mM Tris-HCl pH 7.5, 100 mM NaCl, 10% glycerol, 2 mM DTT, 0.5 mM EDTA), then applied to a Q HP column (Cytiva). After washing with low salt buffer, the protein was eluted using a linear NaCl gradient (100 mM to 1 M) at approximately 460 mM NaCl. Collected fractions were dialyzed overnight against heparin low salt buffer (20 mM Tris-HCl pH 7.5, 50 mM NaCl, 10% glycerol, 2 mM DTT, and 0.5 mM EDTA) and applied to a HiTrap Heparin HP column (Cytiva). Bound protein was washed with low salt buffer and eluted with a linear NaCl gradient (50 mM to 1 M), with the protein eluting at approximately 245 mM NaCl. Pooled fractions were collected and dialyzed twice overnight against storage buffer (20mM Tris-HCl pH 7.5, 50 mM NaCl, 30% glycerol, 2 mM DTT, and 0.5 mM EDTA) and stored at −80°C.

Pellets of clamp loader containing ψ-GS12-χ were resuspended in cell lysis/low salt buffer identical to the ψ-GS8-χ clamp loader purification except that the sodium chloride concentration was lower (20 mM), and clarified as described above. The supernatant was applied to an SP HP column (Cytiva), washed with low salt buffer, and eluted with a linear NaCl gradient (20 to 300 mM) at approximately 140mM NaCl. Pooled fractions were dialyzed overnight against heparin low salt buffer and purified on a HiTrap Heparin column (Cytiva) as for ψ-GS8-χ clamp loader, eluting at approximately 230 mM NaCl. Fractions were dialyzed overnight against Q HP low salt buffer and applied to a Q HP column (Cytiva). After washing with low salt buffer, protein was eluted using a linear NaCl gradient (100 to 500 mM) at approximately 350 mM NaCl. Final pooled fractions were dialyzed twice overnight against storage buffer and stored at −80°C.

Proteins were analyzed by SDS-PAGE under denaturing and reducing conditions using the Bio-Rad Mini-PROTEAN system. Samples were mixed with 6X Laemmli sample buffer containing β-mercaptoethanol and heated at 95°C for 10 min prior to loading. Fusion proteins expressed and purified individually were resolved on 12% polyacrylamide gels.

Gamma complex variants were resolved on 15% polyacrylamide gels to improve the separation of subunits with similar molecular weights. Electrophoresis was performed in standard Tris-glycine-SDS running buffer at 200 V for approximately 60 min for 12% gels and 120 min for 15% gels. Gels were stained with Coomassie Brilliant Blue R-250 and imaged following destaining.

### Differential Scanning Fluorimetry (DSF)

As the pH of Tris buffers varies more with thermal change than HEPES, WT ψχ and each fusion protein, ψ-GS8-χ and ψ-GS12-χ, were dialyzed twice against HEPES-based buffer (20 mM HEPES pH7.5, 50 mM NaCl) to remove Tris from the proteins. Dialyzed protein samples were centrifuged at 11,000 RCF for 5 min at 4°C to remove particulates, and protein concentration measured using a Bradford assay as previously described. The final protein concentration of 0.3 mg mL^-1^ was used for each reaction. Following the manufacturer’s instructions, SYPRO Orange Protein Gel Stain (Invitrogen, Waltham, MA, USA) was diluted 1000X and the melting temperature (T_m_) determined using a temperature range of 10°C to 95°C in increments of 0.5°C every 10 s. A real-time PCR device (CFX Real-Time PCR System, Bio-Rad) was used to monitor protein unfolding by the increase in the fluorescence of the fluorophore SYPRO Orange Assay. Melting temperatures were measured three times (n=3) with technical triplicates in each assay. Statistical analysis was performed in GraphPad Prism using a one-way ANOVA followed by Tukey’s multiple comparisons test (10 comparisons; α = 0.05), and statistically distinct groups were assigned using compact letter display notation.

### Circular Dichroism (CD)

WT ψχ and each elution peak from ψ-χ fusions were dialyzed against buffer containing 10mM KH_2_PO_4_ and 100mM KF to minimize buffer absorption in the far-UV spectra. Proteins were subsequently diluted to a concentration of 0.025 mg mL^-1^. UV CD spectra (185-250 nm) were recorded at 25°C on a Circular Dichroism Spectrometer Model 400 (Biomedical, Inc., Lakewood, NJ, United States). A 1 mm path length cell was used at a step length of 1nm, an averaging time of 5s, a settling time of 0.333s, and a multi-scan wait of 1s. Spectra were corrected by subtracting the dialysis buffer measurements and normalizing by the mean residue weight (MRW) of the proteins, path length (P, cm), and protein concentration (C, mg mL^−1^). Machine units (θ) were obtained in millidegrees from the spectrometer. The final spectra are expressed in Δε (M^-1^ cm^-1^) using Equation 1 (26).

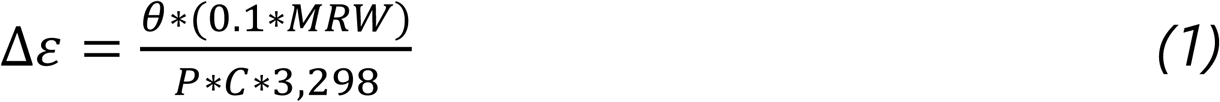

### ATPase-coupled Assay

ATPase activity was measured using a coupled enzyme assay measuring the decrease in NADH absorbance (27), as previously described (14) with slight modifications. Reactions were performed with 0.5μM 20-bp duplex with a dT_65_ 5’ ss overhang (Table 2, SWP1 and YA1comp20T65), 80 nM of γ complex, 1μM of β clamp, 0.75μM of SSB, and 0.5mM of ATP. ATPase activity was measured for WT γ complex, γ complex lacking χ (Δχ), γ complex lacking χ and ψ (min), and γ complex containing the ψ-GS12-χ or ψ-GS8-χ fusion proteins, in the presence or absence of SSB as indicated. The observed rate of ATP hydrolysis was calculated using Equation 2 and normalized to γ complex concentration.

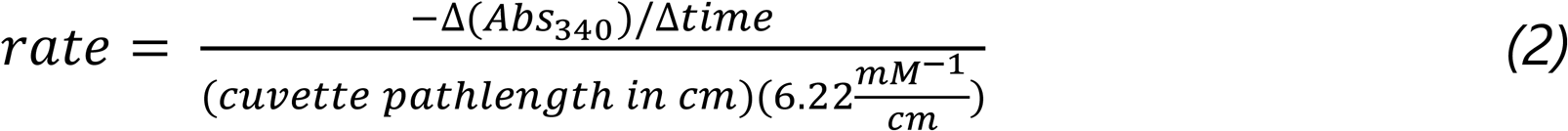

### Labelling β with Fluorophores

Unique surface Cys residues were introduced into β (β-R103C/I305C/C260S/C333S) for clamp closing assays (28). Proteins were labelled as previously described (12, 28). Briefly, Cys residues were labeled with maleimide derivatives for Alexa Fluor 488 (AF488, Invitrogen) or Tetramethylrhodamine (TMR, Invitrogen), creating β-AF488_2_ or β-TMR_2_, respectively, and purified using a desalting column (BioGel P6-DG, BioRad) followed by anion exchange chromatography (HiTrap Q, Cytiva). Labelled β was further dialyzed to remove excess fluorophore. Labeled protein concentrations were determined by using a Bradford-type assay (Bio-Rad Protein Assay) with unlabeled proteins as standards.

### Stopped-flow Clamp Closing Reactions

Reactions were performed according to Newcomb *et al*. with slight modifications. The decrease in AF488 fluorescence accompanying clamp closure on DNA was measured as a function of time using an Applied Photophysics SX20MV stopped-flow instrument. AF488 was excited at 495 nm using a 3.7 nm bandpass, and fluorescence emission was detected using a 515 nm long pass filter. The DNA substrate consisted of a 30-bp duplex and dT_35_ 5’ ss overhang (Table 2, YA1 and JH9T35). A sequential mixing scheme was used in which clamp loader, β-AF488_2_, and ATP were mixed and pre-incubated for 4 s prior to adding DNA and excess unlabeled clamp to measure rates of clamp closing. Final concentrations were 20nM of each clamp loader, 20 nM β-AF488_2_, 200 nM unlabelled β, 80 nM DNA, and 1.5 molar equivalents of SSB per DNA molecule. For WT clamp loader and clamp loaders containing ψ-GS12-χ or ψ-GS8-χ, data were collected every millisecond over four seconds, for a total of 4000 time points. For the clamp loader Δχ, data was collected every millisecond over ten seconds, for a total of 10000 time points. Time courses were fit to double exponential decays (Equation 3) for WT and fusion clamp loaders to estimate the rates of decrease in fluorescence, where *a_fast_* and *a_slow_* are the amplitudes associated with the fast and slow phases, respectively, and *k_fast_* and *k_slow_* are the rates of the fast and slow phases, respectively, and c is a constant. Double exponential decays did not improve the quality of empirical fits of time course reactions with Δχ, so the simpler single exponential fits were used.

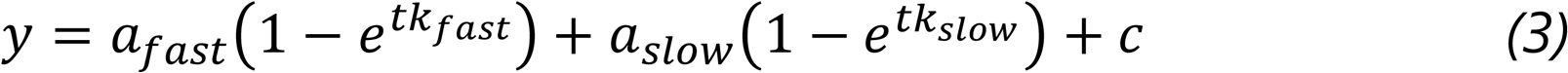

### Fluorescence Measurements in Equilibrium Binding

Reactions were performed as previously described with slight modifications (28). AF488 was excited at 495 nm, emission spectra collected using a 3 nm bandpass, and relative intensity values calculated at 520 nm. Buffer background signals were subtracted from the spectra and relative intensities calculated by dividing intensities for solutions containing γ complex by the intensity for free β. The relative intensity of samples containing no γ complex was set to 1. *K_d_* values were calculated using Equation 4, where β_o_ is the total concentration of γ_o_ is the total concentration of γ complex, and I_max_ and I_min_ are the maximum and minimum intensity, respectively.

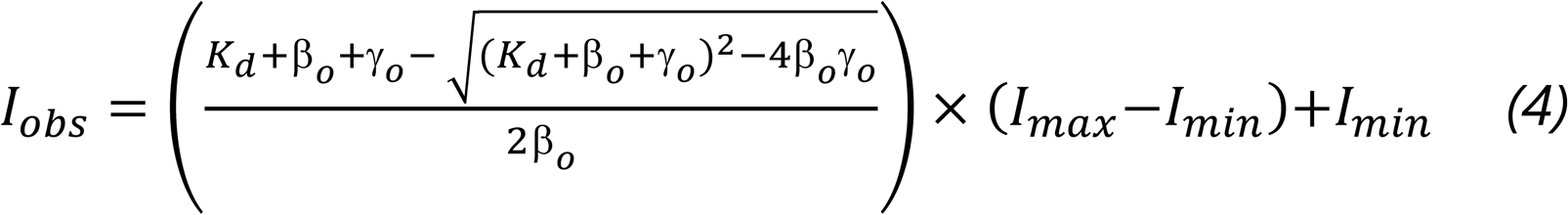

### Stopped-flow Clamp Opening Reactions

The increase in TMR fluorescence accompanying clamp closure on DNA was measured as a function of time using an Applied Photophysics SX20MV stopped-flow instrument. TMR was excited at 540 nm using a 5.6 nm bandpass, and fluorescence emission was detected using a 570 nm long pass filter. A single mixing scheme was used in which β-TMR_2_ was mixed with variations of clamp loader. Final concentrations were 60nM of each clamp loader, 40 nM β-TMR_2_, and 0.5 mM ATPγS. Data was collected every millisecond over ten seconds, for a total of 10000 time points. Time courses were fit to double exponential rises (Equation 5).

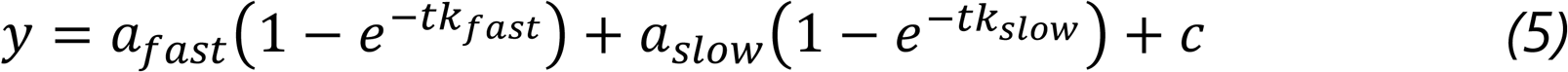

### Yeast Two-hybrid Analysis

Yeast two-hybrid assays were performed as previously described, with slight modifications, using the Matchmaker™ GAL4 Two-Hybrid System 3 (Takara Bio)(19). Full-length *E. coli yoaA* (wild-type or T620A mutant), *ssb*, *holC*, and a *holD*-*GS12*-*holC* fusion were cloned into vectors encoding either the GAL4 DNA-binding domain (BD) or GAL4 activation domain (AD), as indicated. Negative controls included empty BD or AD vectors paired with each hybrid construct, as well as combinations recommended by the Matchmaker system to assess background reporter activation. In addition, each BD and AD construct was tested individually to assess autoactivation of the GAL4 reporter system, and no autoactivation was observed under the conditions used. Single colonies of yeast carrying the appropriate plasmid pairs were grown overnight (∼20 h) at 30 °C in selective dropout medium lacking leucine (-LEU) or tryptophan (-TRP) to maintain plasmid selection. Cultures were diluted in sterile water prior to spotting to normalize cell density. Strains expressing SSB were diluted 1:15, while all other strains were diluted 1:25. Equal volumes of diluted cultures were spotted onto synthetic dropout plates lacking leucine (-LEU), tryptophan (-TRP), or histidine (-HIS). Growth on -LEU and -TRP plates confirmed retention of the respective BD and AD plasmids, while growth on -HIS plates was used to assess activation of the HIS3 reporter gene as an indicator of protein-protein interaction. Plates were incubated at 30 °C for 2-3 days prior to imaging.

### Western Blot Analysis

Protein expression under complementation conditions was evaluated by immunoblotting in WT *E. coli* cells to obtain sufficient cell mass for fractionation. The WT parent strain BW25113 was streak-isolated prior to preparing chemically competent cells. Cells were grown under conditions identical to those used for complementation and survival assays below.

WT strains were transformed with pBAD18 plasmids containing the empty vector, pBAD18_*holC*, pBAD18_*holD*, pBAD18_*holD*_*holC*_operon, pBAD18_*holD*-GS12-*holC*, pBAD18_*holD*-GS12-*holC*_R128A, and plated on LB + 0.2% glucose. Colony isolates were inoculated into liquid LB + 0.2% glucose and grown overnight at 37°C without shaking. Cultures were normalized to OD_600_ = 0.05 in the same media and grown for 2 h at 37°C, with shaking at 250 rpm. Five replicates were plated at a 10^-3^ dilution on LB agar with glucose or arabinose. Cells were harvested with an inoculation loop, pooled, and resuspended in 1 mL of Phosphate-Buffered Saline (PBS), then centrifuged at 13,000 rpm for 1 min. The pellets were weighed and stored at −20 °C until processing.

Cell pellets were lysed using NEBExpress *E. coli* Lysis Reagent (New England Biolabs) at 5 mL g^-1^ of cell mass and incubated at room temperature for 20 min, with inversions every 2 minutes. Lysates were centrifuged at 13,000 rpm for 10 min at 4 °C to separate supernatant and pellet fractions. Protein concentration in supernatant fractions was measured at 280 nm using a NanoDrop spectrophotometer and normalized to 5.3 mg mL^-1^ using lysis reagent and Laemmli buffer. Pellet fractions were resuspended in 5 μL mg^-1^ of 6x Laemmli sample buffer. All samples were heated at 95 °C for 10 min prior to electrophoresis.

For each condition, 2 μg of total protein from the soluble fraction was loaded per lane, along with an equivalent volume of the corresponding pellet fraction. Equal protein loading was ensured by normalization of total protein concentration. Proteins were resolved on 15% SDS-PAGE gels and transferred to 0.2 μm PVDF membranes at 55 V for 50 min at 4 °C. Identical samples were run on separate gels to detect χ, ψ, and ψ-GS12-χ in supernatant and pellet fractions.

All washes included Tris-Buffered Saline with Tween-20 (TBST) for 5, 5, and 15 min. Membranes were washed and subsequently blocked for 1 h at room temperature in 5% (w/v) nonfat milk in TBST. Following another round of washes, membranes were incubated overnight at 4 °C with primary antibodies diluted 1:5,000 in 5% milk TBST. After additional washes, membranes were incubated with HRP-conjugated secondary antibodies diluted 1:20,000 in 5% milk TBST for 1 h at room temperature, then washed again.

For detection, blots were developed using 2 mL of Pierce™ ECL Western Blotting Substrate (Thermo Scientific) and imaged on a Bio-Rad ChemiDoc (Bio-Rad Laboratories) system with automatic optimal exposure settings. The images were then inverted and auto-scaled using the Image Lab Software for ChemiDoc.

Chi was detected with a rabbit polyclonal antibody, and ψ with a mouse monoclonal antibody, both provided by Charles McHenry’s laboratory. The secondary antibodies used were goat anti-rabbit IgG-HRP and goat anti-mouse IgG-HRP (Bio-Rad Laboratories).

### Complementation and Survival Assays

*E*. *coli* KEIO *holC* knockout strain JW4216 and WT *E. coli* were streak-isolated prior to preparing chemically competent cells to confirm the expected phenotype, including reduced growth and small colony size, and transformants used in the experiments were verified to display the correct colony phenotype. These strains were then transformed with pBAD18 plasmids containing the empty vector, pBAD18_*holC*, pBAD18_*holD*, pBAD18_*holD*_*holC*_operon, pBAD18_*holD*-GS12-*holC*, pBAD18_*holD*-GS12-*holC*_R128A, and plated on LB + 0.2% glucose. Colony isolates were inoculated into liquid LB + 0.2% glucose and grown overnight at 37°C without shaking. Cultures were normalized to OD_600_ = 0.05 in the same media and grown for 2 h at 37°C, with shaking at 250 rpm. Cells were then washed in LB, serially diluted, and spotted using 4 µL onto LB agar plates containing Carb100, either 0.2% glucose or 0.2% arabinose, and with or without AZT at 12.5 ng mL^-1^ or 25 ng mL^-1^. Plates were incubated overnight at 37°C. Colony counts and all calculations are provided in the Supplemental Data file.

### Buffers

Reaction buffers contained 20mM Tris-HCl pH 7.5, 50 mM NaCl, 8 mM MgCl_2_, 0.5 mM ATP, 5 mM DTT, 0.5mM EDTA, 40 μg mL^-1^ BSA, and 4% glycerol.

### Reproducibility and Error

Experiments were generally done three times. As an additional measure of robustness, measurements were made by comparing different reactions side-by-side (kinetic) to rule out systematic effects due to methods. Representative kinetic traces are shown. Rate and equilibrium constants were calculated from three independent experiments and are reported as the average with standard deviation. Titration data for each of the three replicate experiments are plotted along with the average and standard deviation.

## RESULTS

### Design of ψ-χ Fusions

Two ψ-χ fusion proteins were designed based on the solved heterodimer structures (PDB: IEM8 (29), 3SXU (30)). The distance between the C-terminus of ψ and the N-terminus of χ was measured using ChimeraX (∼30Å). To account for missing residues within the structures (one in 1EM8 and four in 3SXU) and to facilitate accurate linker spacing within the globular protein structure, two linkers were designed: GS12 (12 residues) and GS8 (8 residues). Each consisted of glycine-serine repeats (Table 2), chosen for their flexibility and hydrophilicity, which promote solubility and reduce steric hindrance, as described in prior studies (31–34).

Using AlphaFold Colab (20), models of ψ-GS8-χ and ψ-GS12-χ predicted that both GS8 and GS12 linkers were unstructured and of sufficient length to span the region between the C-terminus of ψ and N-terminus of χ (Figure 2). Each ψ-χ fusion sequence was synthesized to ensure the rigor and viability of the fusion proteins in vitro.

**Figure 2:**
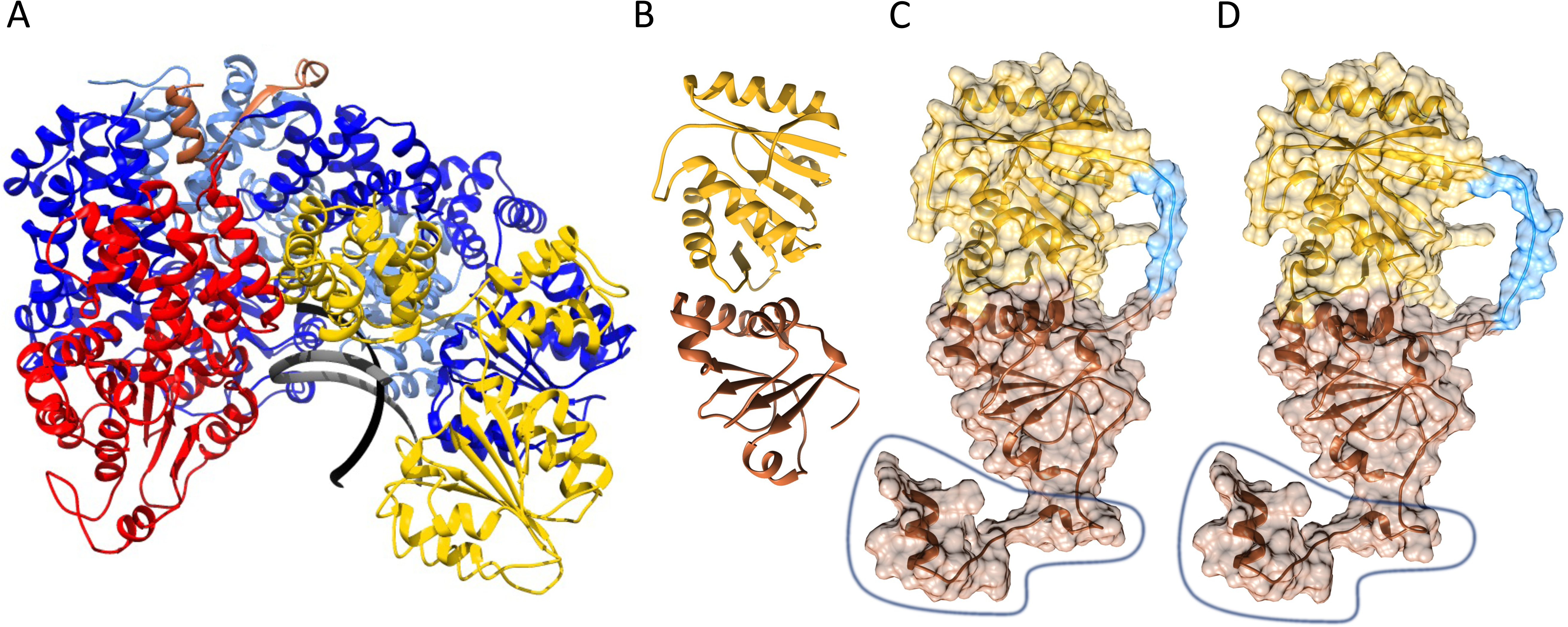
Ribbon diagrams of the WT clamp loader complex and ψ-χ fusion proteins. A) The clamp loader (PDB ID: 3GLI) (γ_3_δδ′, blue, yellow, and red, respectively) is bound to a peptide derived from the N-terminal of ψ (brown). The DNA primer and template strand are shown in grey. B) WT ψχ (PDB ID: 3SXU) structure (ψ in brown and χ in gold), has 33 unmodeled residues in the N-terminal of ψ and 4 unmodeled residues in the N-terminal of χ. Fusion models made with AlphaFold 2 Colab for C) ψ-GS8-χ and D) ψ-GS12-χ have linkers shown in light blue. The circled regions in C and D represent the peptide of ψ present in the clamp loader crystal structure in A.

### ψ-GS12-χ is More Stable Than ψ-GS8-χ

To biochemically characterize the fusion proteins independently of the clamp loader, each fusion protein was expressed with an N-terminal 6X-His tag and purified by nickel affinity chromatography followed by cation-exchange chromatography (Figure 3, Supplemental Figure 1). The ψ-GS12-χ fusion was successfully purified, yielding two distinct peaks on the cation exchange column. In contrast, ψ-GS8-χ eluted as two peaks from the nickel column (Supplemental Figure 2), but was insufficiently soluble for further purification by cation exchange.

**Figure 3:**
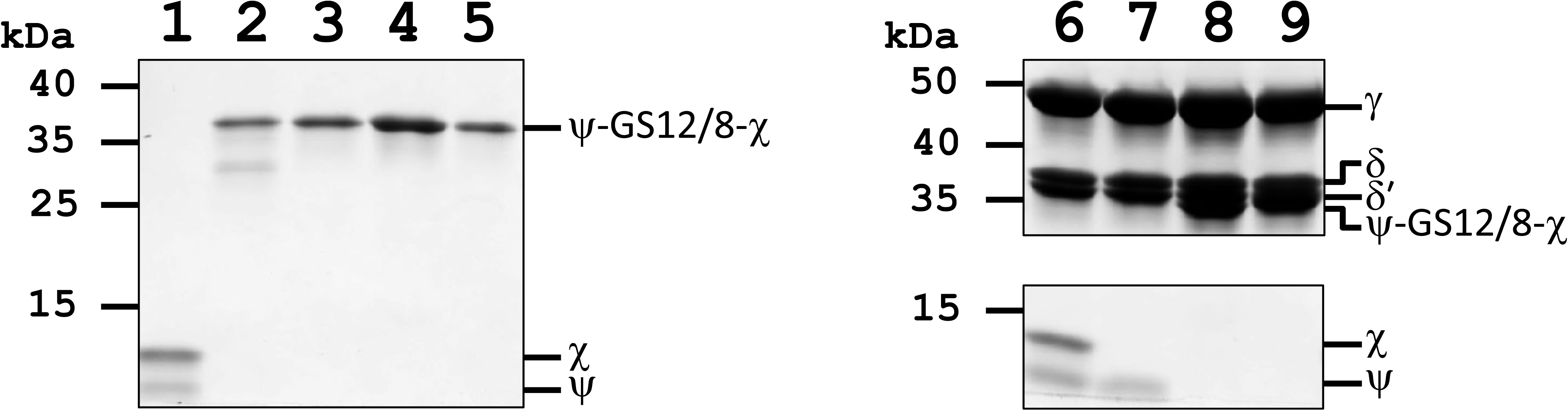
SDS-PAGE analysis of purified ψ-χ fusion proteins and γ complex variants used in this study. Lane 1, ψχ; lane 2, ψ-GS12-χ #1; lane 3, ψ-GS12-χ #2; lane 4, ψ-GS8-χ #1; lane 5, ψ-GS8-χ #2; lane 6, WT γ complex; lane 7, γ complex Δχ; lane 8, γ complex ψ-GS12-χ; lane 9, γ complex ψ-GS8-χ. Proteins were resolved by SDS-PAGE under denaturing conditions and stained with Coomassie Brilliant Blue. Major bands migrate at the expected molecular weights for the indicated complexes and fusion proteins. All preparations were used for the biochemical and kinetic analyses described in the text. The uncropped gel is provided in Supplemental Figure 3.

The thermal stability of the fusion proteins was assessed using differential scanning fluorimetry (DSF) using SYPRO Orange in HEPES buffer (Figure 4A-B). Both ψ-GS12-χ fusion peaks exhibited melting temperatures (T_m_) approximately 5 ℃ higher than WT ψχ (Figure 4A-B). For ψ-GS8-χ, the first and second purification fractions had a T_m_ of approximately 1.5 ℃ higher and 2 ℃ lower, respectively. The ψ-GS8-χ fusion also exhibited greater variability in the T_m_ for the second peak, possibly indicating variations in protein folding or changes in the oligomerization state of the protein.

**Figure 4:**
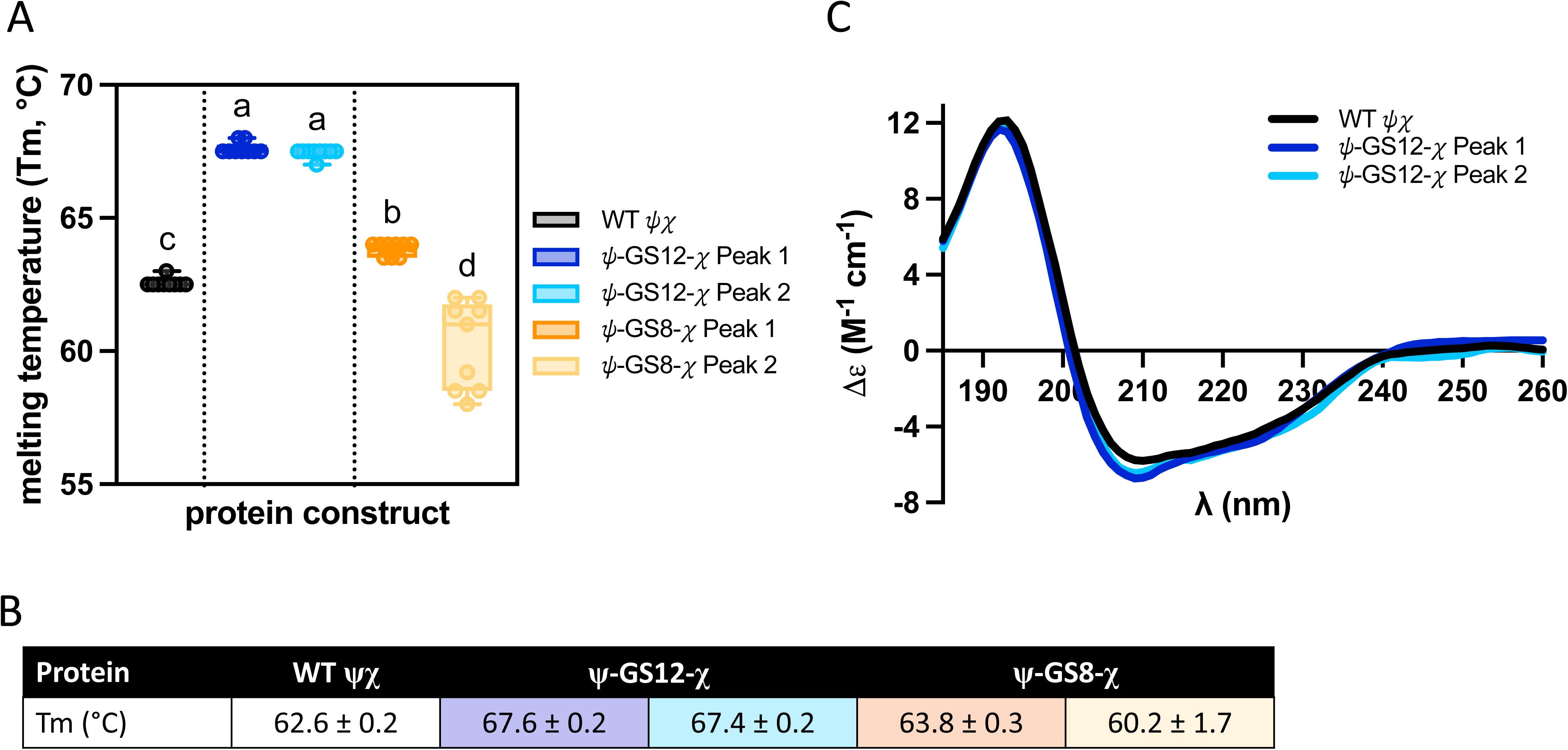
Modifications to ψχ and their effects on protein stability. A-B) Melting temperature (T_m_) of WT ψχ and each fusion protein, ψ-GS12-χ and ψ-GS8-χ, was determined using SYPRO Orange and a temperature range of 10°C to 95°C in increments of 0.5°C every 10 s. T_m_ values were compared by one-way ANOVA with Tukey’s multiple comparisons test (familywise α = 0.05, 10 comparisons). Compact letter display labels above each bar indicate statistically distinct groups; conditions that do not share a letter differ significantly (P < 0.05). B) Averages and standard deviation of T_m_’s reported in panel A. C) Purified WT ψχ (black) and ψ-GS12-χ peak one (dark blue) and two (light blue) analysed by far-UV circular dichroism spectra. Each curve is an average of three scans, normalized by the mean residue weight of each protein, path length, and protein concentration.

Circular dichroism (CD) was used to assess whether the fusions fold the same as WT. The CD spectra of each ψ-GS12-χ peak (Figure 4C) were nearly identical to one another and highly similar to that of WT ψχ, suggesting the fusion does not alter folding. The ψ-GS8-χ fusion precipitated under CD buffer conditions, preventing analysis. Collectively, these results demonstrate that the ψ-χ fusions are stable according to DSF, and the CD spectrum of ψ-GS12-χ closely resembles that of WT ψχ. However, the ψ-GS8-χ tends to be less soluble.

### Ψ-χ Fusions are Biochemically Active in the Context of the γ Complex Clamp Loader

While the individual ψ-χ fusions were stable, evaluating their functionality within the clamp loader is crucial. To test this, ψ-χ fusions proteins were co-expressed with γ complex subunits to form γ complexes containing either ψ-GS12-χ (γ_3_δδ’ψχ-GS12-χ) or ψ-GS8-χ (γ_3_δδ’ψχ-GS8-χ) without N-terminal his-tags (Figure 3, Supplemental Figure 1).

Clamp loader activity was validated by comparing fusion-containing complexes to WT γ complex across several stages of the clamp loading process. The γ complex functions as a circular pentamer, where ATP binding by the γ-subunits drives conformation changes that promote β-clamp binding and subsequent DNA loading. ATP hydrolysis then triggers clamp release. ATP hydrolysis is a critical step in the clamp opening cycle, driving the conformational transitions required for both opening and releasing the β-clamp on DNA. Clamp loaders that lack ψ are defective in specific ATP-dependent steps, affecting both the efficiency and temporal order of clamp loading (35). Therefore, verifying ATP hydrolysis activity in the fusion-containing complexes is essential to establish that the fusions functionally integrate into the clamp loader. ATP hydrolysis was measured using an enzyme-coupled assay that links ADP production to NADH oxidation (27). In addition to the γ complexes containing the ψ-χ fusion proteins, ATPase activity was measured for γ complex lacking χ, as well as a minimal complex lacking both ψ and χ, in the presence and absence of SSB. Under these conditions, all γ complex variants exhibited comparable steady-state ATP hydrolysis rates, with observed rates normalized to γ complex concentration ranging from approximately 4-6 s^-1^ (Supplemental Figure 3). The inclusion of SSB did not alter ATP activity in any of the tested conditions. Thus, incorporation of the ψ-χ fusions did not impair the ATPase activity of the clamp loader in steady-state assays.

Steady-state ATPase rates reflect the rate-limiting step in the complete catalytic cycle for clamp loading which may not be sensitive to χ-SSB interactions. In contrast to steady-state ATPase reactions, χ plays a critical role during the pre-steady state clamp loading at SSB-coated ss/ds DNA junctions (12). In our stopped-flow assay monitoring β-clamp closing by fluorescence quenching, SSB inhibits the clamp-closing step in the absence of χ, demonstrating that the pre-steady state rate of clamp closing depends on χ-SSB interactions. Fusion-containing complexes were therefore assessed for clamp-closing activity on DNA-SSB substrates (Figure 5A).

**Figure 5:**
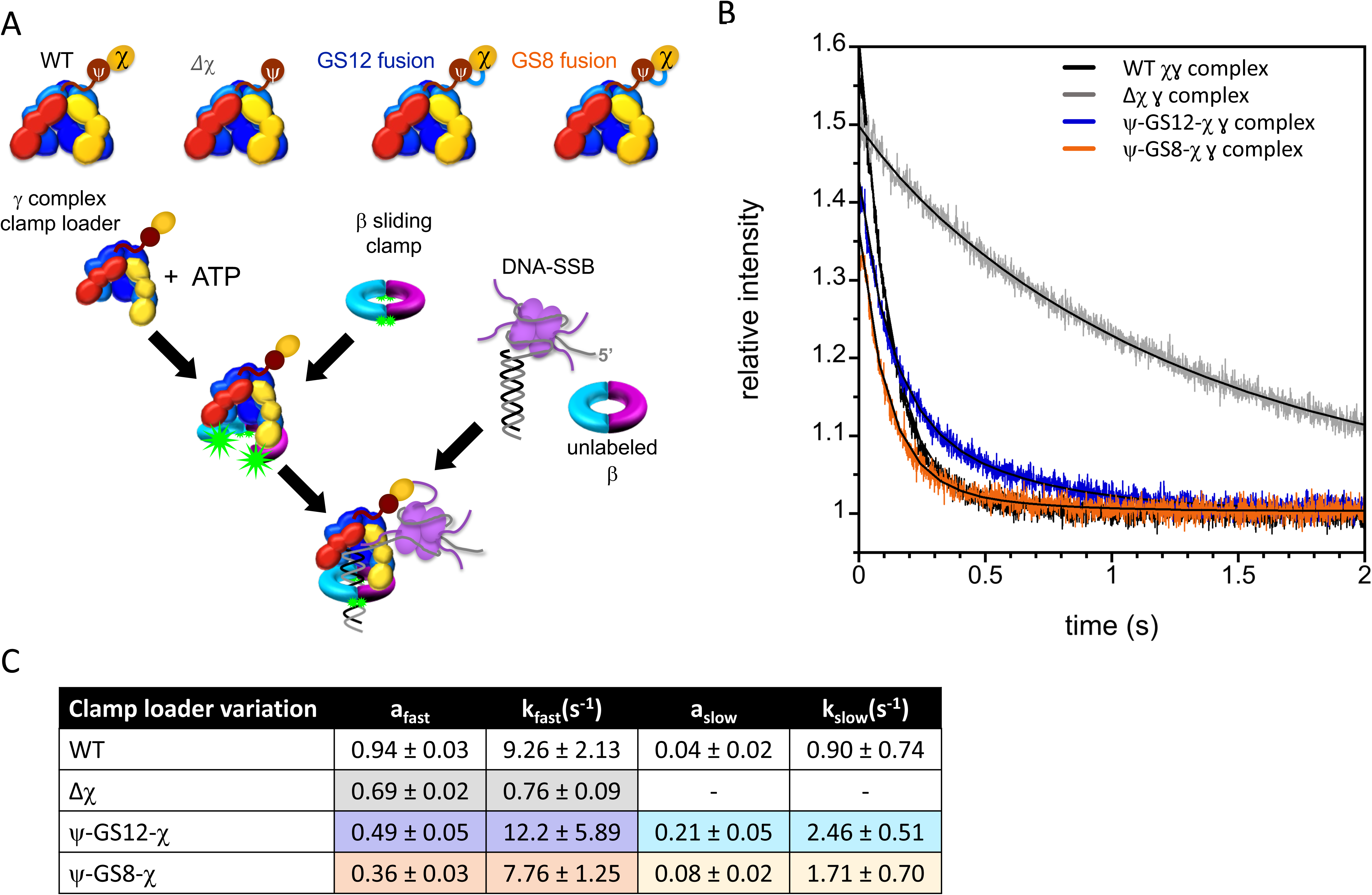
Fusion clamp loading onto DNA. A) Clamp loading on DNA with bound SSB. The reaction scheme for clamp closing is shown alongside γ complex variations. The clamp loader was pre-incubated with β-AF488_2_ and ATP for 4 s to form an open clamp loader-clamp complex prior to adding DNA-SSB along with an excess of unlabeled β-clamp to limit reactions to a single turnover. B) Representative clamp closing reactions for containing DNA-SSB along with 20 nM γ complex clamp loader, 20 nM β-AF88_2_, 200 nM unlabeled β, 80 nM DNA, and 1.5 molar equivalents of SSB per DNA molecule, with 0.5 mM ATP. Gamma complex variations are shown as WT (black), Δχ (grey), ψ-GS12-χ (blue), and ψ-GS8-χ (orange). Solid black lines through the data shown fit to a double exponential (Equation 3) with values in Figure 5B. C) Rates from the kinetic fits as described in Methods.

The β sliding clamp was labeled with Alexa Fluor 488 (AF488) at both sides of the dimer interface (12, 28). In this system, when β-AF488_2_ closes on DNA, the fluorophores are brought within quenching distance, resulting in low relative fluorescence (12, 28, 36).

Conversely, when β-AF488_2_ is open and bound to the γ complex, the fluorophores are separated, leading to high relative fluorescence. A DNA substrate consisting of a 30-bp duplex and dT_35_ 5’ ss overhang was used. Clamp-closing rates of WT γ complex were compared to those of the γ complex lacking χ (γ_3_δδ’ψ) and complexes containing either fusion (ψ-GS12-χ or ψ-GS8-χ). Rates were measured using a sequential-mixing stopped-flow experiment in which γ complex, β-clamp, and ATP were mixed and pre-incubated (4s) to form an open clamp loader-clamp complex before the addition of DNA-SSB. An excess of unlabeled β-clamp was also added to limit reactions to a measurable single-turnover event (12). While time courses for clamp closing are not exponential, data were fit to exponential decay to estimate the rate of decrease in fluorescence and measurements of reproducibility.

As expected, in the absence of χ (Δχ), γ complex was inhibited by SSB, with clamp loading occurring approximately 5-fold more slowly than WT (Figure 5B-C). In contrast, γ complexes containing either ψ-χ fusion closed clamps at rates indistinguishable from within experimental error (Figure 5C), demonstrating that both fusion proteins functionally interact with SSB to remodel DNA-SSB and clamp loading. The amplitudes of clamp closing reactions were reduced for the fusion complexes compared to WT, with γ complex containing GS8 showing the lowest amplitude.

To determine whether these amplitude differences arose from altered clamp-loader affinities, dissociation constants (*K_d_*) for β-AF488_2_ (20 nM) were measured in the presence of ATP (0.5 mM) (Figure 6A) (28). WT γ complex bound β-AF488_2_ with a *K_d_*of 2.0 ± 0.2 nM, consistent with previous values for β (3.2 nM, (37)) and β-AF488_2_ (3.3 ± 0.2 nM, (28)). The ψ-GS12-χ γ complex exhibited a similar *K_d_* of 2.1 ± 0.3 nM, while ψ-GS8-χ was 4.2 ± 2.7 nM (Figure 6B-C). Thus, clamp loader-clamp binding affinities are the same within error for all three clamp loaders. At the concentrations used in stopped-flow clamp closing experiments, the percent of clamp loader-clamp complexes present when DNA is added is 73% for WT and ψ-GS12-χ clamp loader and 63% for ψ-GS8-χ.

**Figure 6:**
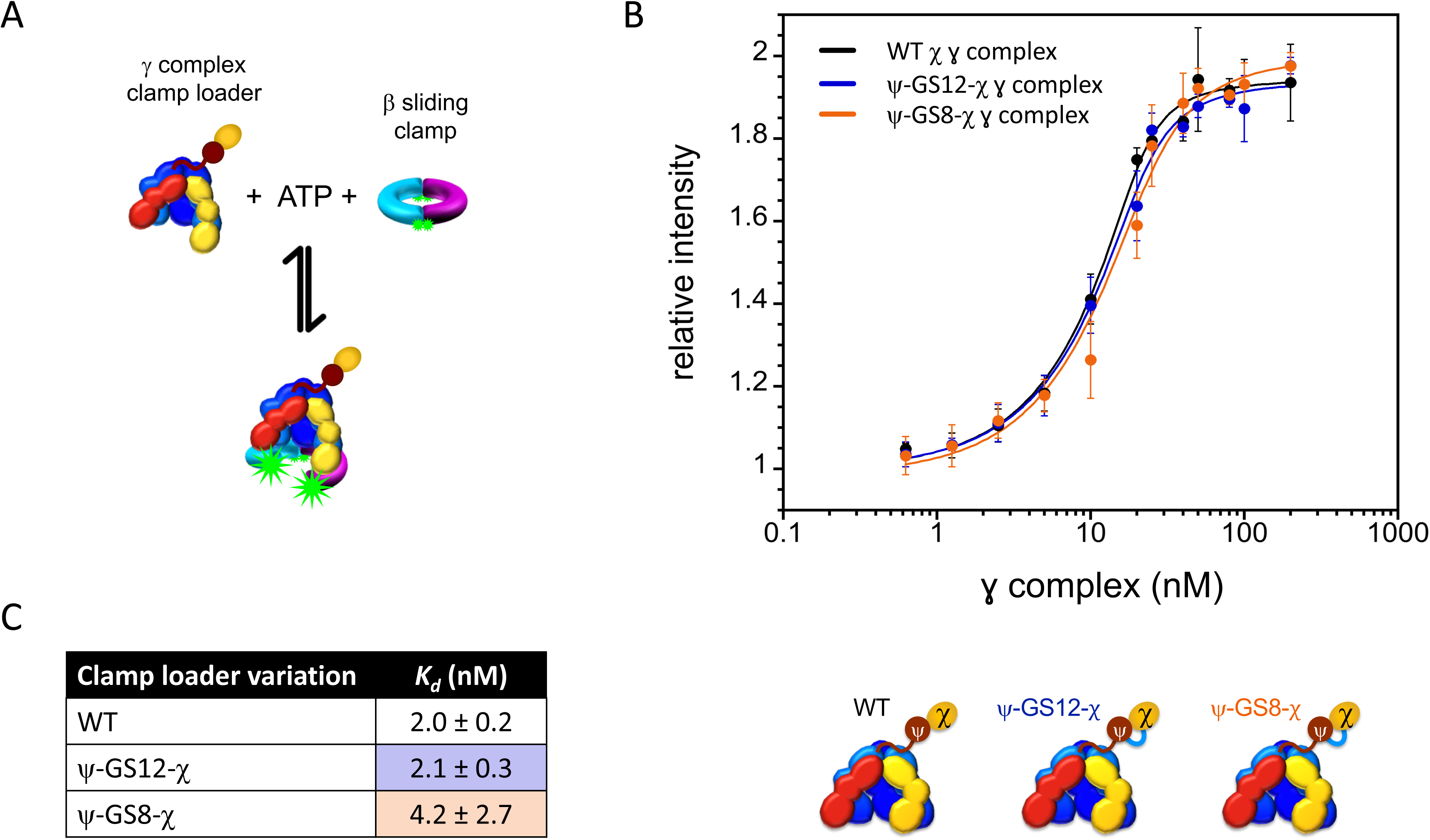
Equilibrium binding of clamp loader fusions to β. A) The reaction scheme for clamp opening is shown using WT γ complex the clamp loader is incubated with β-AF488_2_ and ATP to form an open clamp loader-clamp complex. B) The relative intensity of AF488 at 520 nm is plotted as a function of γ complex concentration for solutions containing 10 nM β-AF488_2_ and 0.5 mM ATP. Gamma complex variations are shown as WT (black), ψ-GS12-χ (blue), and ψ-GS8-χ (orange). C) Data from individual experiments were fit to Equation 4 to calculate the average dissociation constants.

The maximal fluorescence intensities at saturation were equivalent, indicating that the same fraction of clamps was open in all cases. Thus, the reduced amplitudes observed for the ψ-χ fusions cannot be explained by differences in clamp opening defects and are at least partly attributable to the lower fraction of bound clamp loader-clamp complexes for ψ-GS8-χ.

Amplitude differences in clamp closing reactions may also likely reflect subtle changes in clamp loader-clamp dynamics. To test whether clamp opening kinetics contributed to these differences, opening reactions were monitored during the pre-incubation step. Both ψ-χ fusion γ complexes opened clamps comparable to WT, and the four-second pre-incubation was sufficient for complete clamp opening (Supplemental Figure 4).

### Tethering χ to ψ Disrupts Its Interaction with YoaA While Preserving SSB Binding

Chi engages ψ and YoaA through overlapping interaction surfaces, such that incorporation of χ into the clamp loader via interactions with ψ is incompatible with simultaneous YoaA binding (13). The ψ-χ fusion proteins retain replication-associated activity within the clamp loader, and this design constraint predicts that tethering χ to ψ should eliminate its interaction with YoaA while maintaining interactions with SSB. To directly test this, we performed yeast two-hybrid (Y2H) analysis.

Consistent with previous studies, WT YoaA produced a robust interaction with χ, as indicated by growth on the histidine-deficient (-HIS) medium, whereas the YoaA_T620A mutation abolished this interaction in both fusion orientations (19), confirming assay specificity (Figure 7). Chi also exhibited a strong interaction with SSB, as expected. In contrast, no detectable interaction between YoaA and the ψ-GS12-χ fusion was observed in either domain orientation. Importantly, the ψ-GS12-χ fusion retained clear interaction with SSB in the same assay (Figure 7).

**Figure 7:**
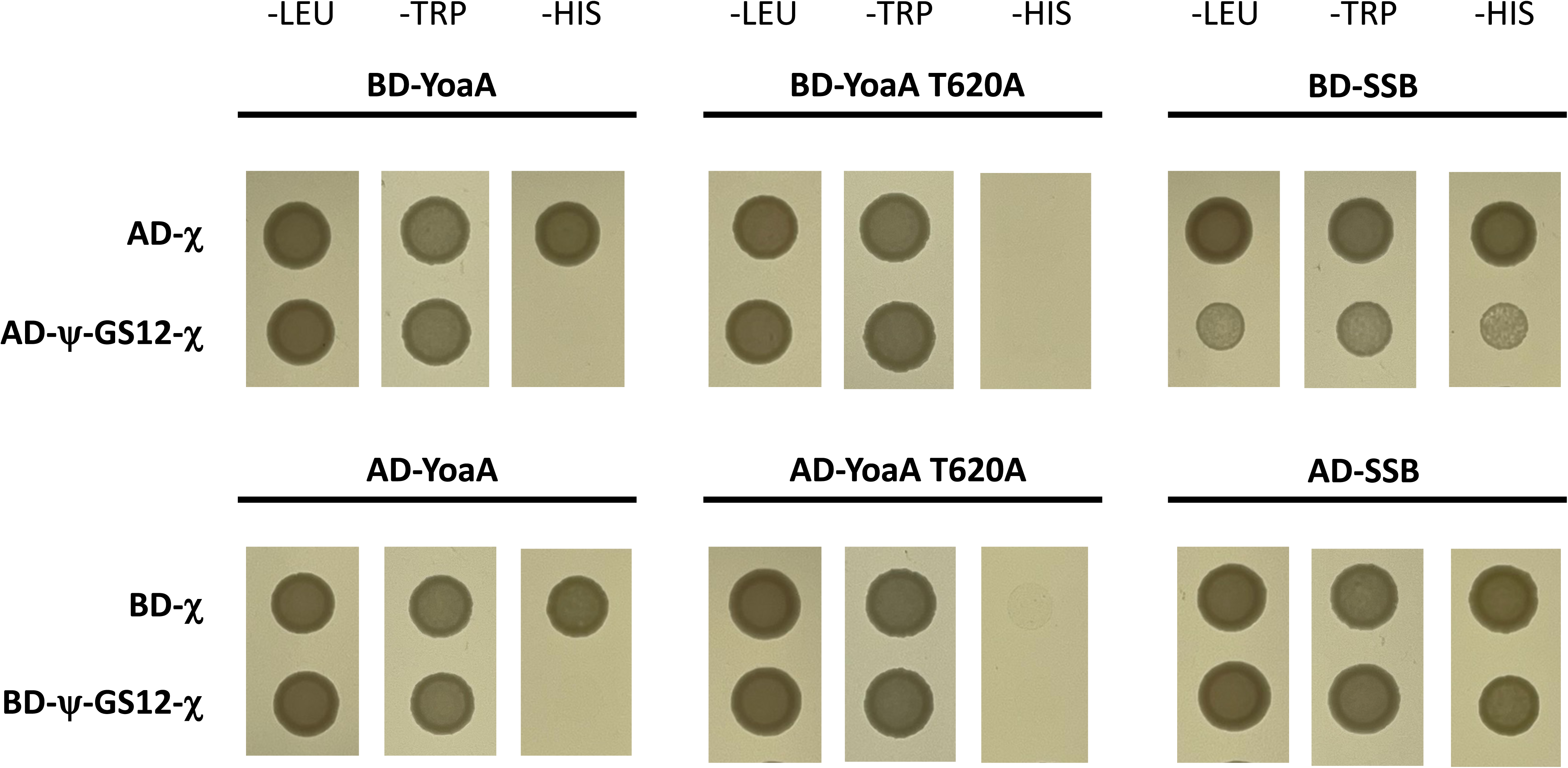
Yeast two-hybrid analysis of interactions among YoaA, SSB, χ, and the ψ-GS12-χ fusion. Yeast two-hybrid assays were performed using GAL4 DNA-binding domain (BD) and activation domain (AD) fusions. Growth on selective dropout medium lacking leucine (-LEU) and tryptophan (-TRP) plates confirms retention of the BD and AD plasmids, respectively, while growth on medium lacking histidine (-HIS) plates indicates activation of the HIS3 reporter and a detectable protein-protein interaction. Top panels: YoaA (wild-type or T620A χ-interaction-defective mutant) or SSB fused to the BD, paired with χ or the ψ-GS12-χ fusion fused to the AD. Bottom panels: Reciprocal orientation, with YoaA or SSB fused to the AD and χ or the ψ-GS12-χ fusion fused to the BD. Strains expressing SSB were diluted 1:15, while all other strains were diluted 1:25 prior to spotting to maintain comparable cell density across conditions.

These data demonstrate that tethering χ to ψ selectively eliminates detectable YoaA interaction while preserving SSB binding. When considered alongside the replication competence of the fusion protein, these results indicate that the ψ-GS12-χ fusion should function within the DNA polymerase III holoenzyme in vivo but cannot engage the YoaA helicase.

### Neither WT ψ-χ or ψ-GS12-χ Rescues AZT Sensitivity of *E. coli* Lacking χ

The fusions demonstrated functionality in vitro, and the next objective was to assess their ability to function in vivo. Both fusion proteins function within the context of the clamp loader, but the ψ-GS8-χ fusion protein is generally less soluble, which precluded some measurements. Therefore, in vivo measurements were done with the ψ-GS12-χ fusion only.

To address the function of the fusion in cells, Δ*holC* and WT *E. coli* strains were transformed with plasmids encoding *holD*-GS12-*holC* under the control of the arabinose-inducible pBAD promoter. Additional control plasmids enabled the expression of *holD* (ψ) alone, *holC* (χ) alone, or both *holD* and *holC* in tandem (*holD*-*holC*_operon), from the same vector backbone.

Expression under induction conditions was confirmed by immunoblotting (Figure 8, Supplemental Figure 5). WT cells were used to obtain sufficient material, although induction conditions were identical in WT and Δ*holC* strains. Endogenous χ was detected in all strains, consistent with its high cellular abundance (38, 39). Upon arabinose induction, χ expressed alone or co-expressed with ψ was detected predominantly in the soluble fraction. Expression of ψ alone yielded protein in the pellet with no soluble signal, consistent with previous reports that ψ requires stoichiometric amounts of χ for solubility (40). The ψ-GS12-χ fusion and R128A variant were detected in the soluble fractions at the expected higher molecular weight, while the R128A variant was also detected in the pellet fraction. These data confirm the expression of proteins tested in the phenotypic and AZT survival assays.

**Figure 8:**
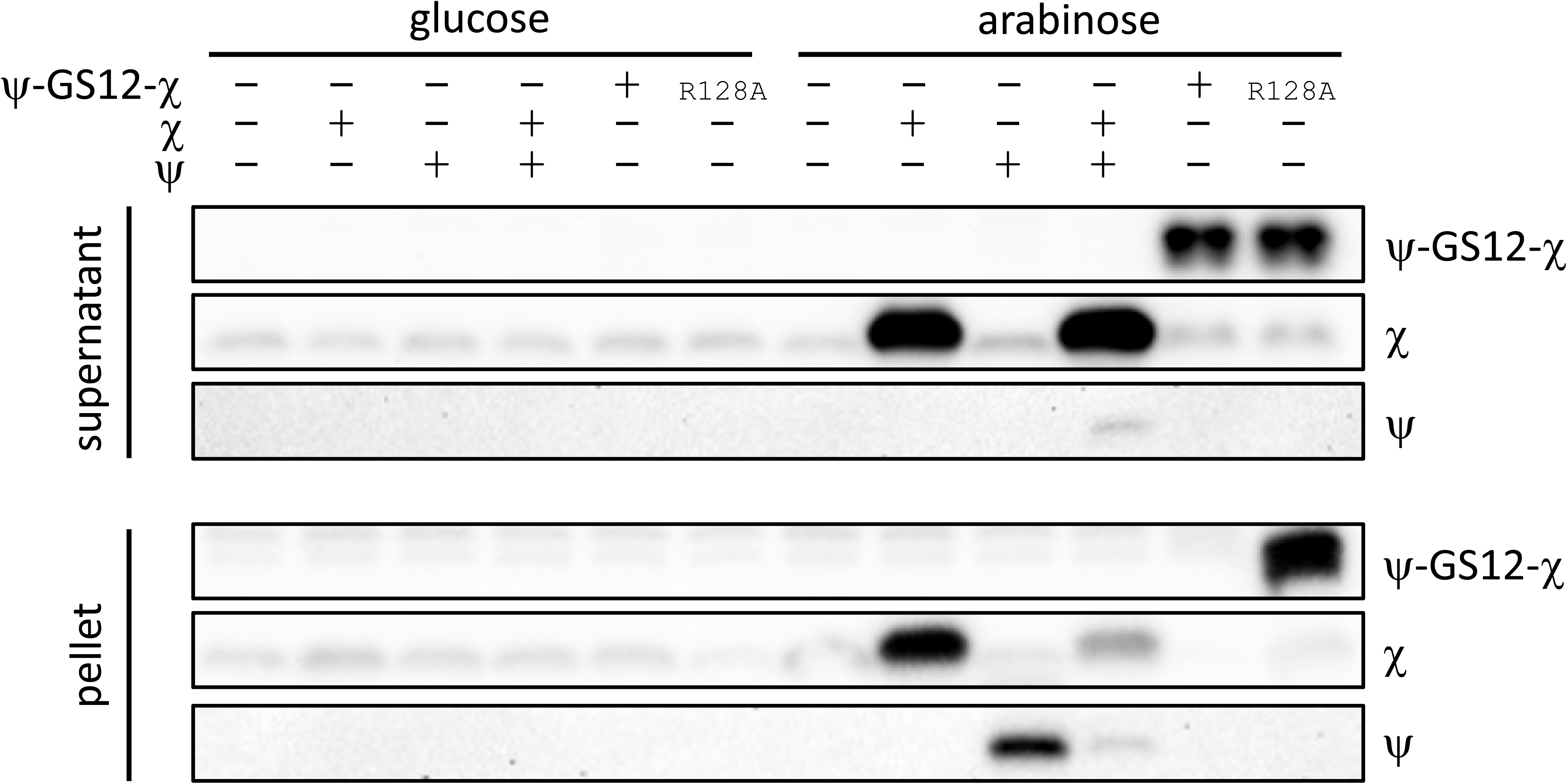
Representative Western blot confirming expression of χ, ψ, and the ψ-GS12-χ fusion constructs under complementation conditions. WT *E. coli* cells were grown under the same induction conditions used for complementation assays, lysed, and fractionated into soluble (supernatant) and insoluble (pellet) fractions. Equal amounts of total protein were loaded per lane based on A280 quantification. Blots were probed with antibodies against χ or ψ, as indicated. A bar above the blot indicates carbon source, with the first six lanes grown in glucose (repressed conditions) and the second six lanes grown in arabinose (induced conditions). Plus (+) and minus (−) symbols above the blot denote the presence or absence of expression of χ, ψ, or the ψ-χ fusion in each lane.

Strains lacking χ (Δ*holC*) are hypersensitive to AZT compared with WT strains, yet it is unclear whether χ contributes to tolerance through its role in the pol III HE, through its interaction with the YoaA helicase, or through a combination of these activities(17). To distinguish these possibilities, AZT sensitivity was measured in *ΔholC* and WT backgrounds expressing plasmid-borne genes (Figure 9). Panels A and D show fractional survival at increasing AZT concentrations for *ΔholC* (Figure 9A) and WT (Figure 9D) strains. For each strain, colony counts at each AZT concentration were normalized to the no-AZT condition, allowing comparison of relative AZT sensitivity independent of baseline growth, with calculations provided in Supplemental Data. Panels B and E show the corresponding total viable cell counts in the absence of AZT for *ΔholC* (Figure 9B) and WT (Figure 9E), providing a measure of baseline colony-forming units. Panels C and F show representative serial dilution plates corresponding to the quantified data.

**Figure 9:**
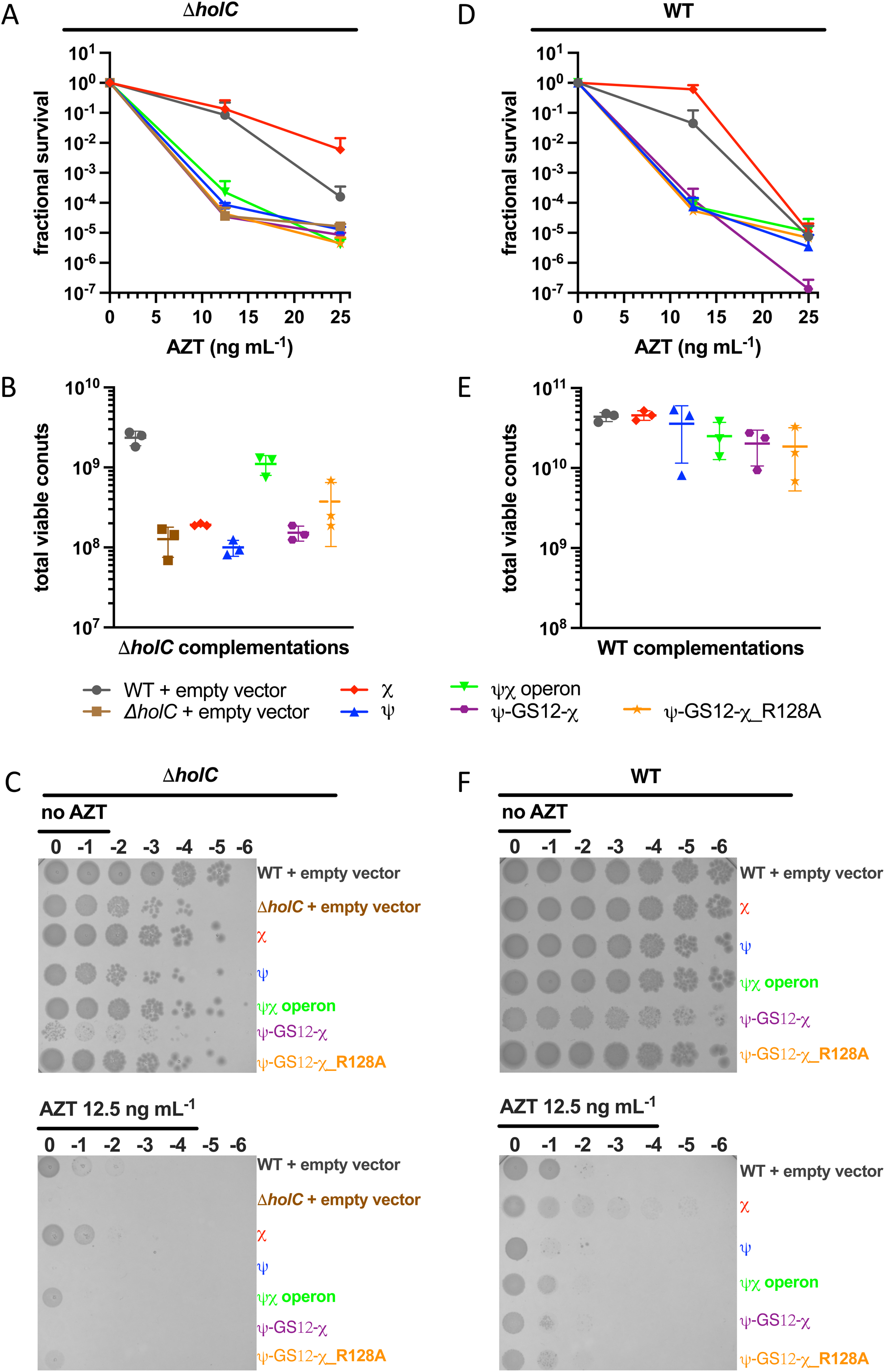
Plasmid-based complementation of ψ-GS12-χ in Δ*holC* (left panels A-C) and WT *E. coli* (right panels D-F). Empty vector controls are shown in Δ*holC* cells (brown squares) and WT cells (grey circles). Both Δ*holC* and WT cells were complemented with plasmids expressing *holC* (χ, red diamonds), *holD* (ψ, blue upward triangles), *holD*-*holC* operon (ψχ, green inverted triangles), *holD*-GS12-*holC* (ψ-GS12-χ, purple hexagons), and *holD*-GS12-*holC*_R128A (ψ-GS12-χ_R128A, orange stars). A and D) Fractional survival following AZT treatment. B and D) Total viable cell counts measured in the absence of AZT (AZT = 0), representing the colony-forming units used as the reference condition for survival normalization. C and F) Representative ten-fold serial dilution spot assays plated on LB medium with or without AZT in the presence of 0.2% arabinose. All constructs were expressed from pBAD18-derived plasmids under arabinose induction, with “Δ*holC* + empty vector” and “WT + empty vector” referring to the empty pBAD18 control in the respective strain backgrounds. Survival was calculated relative to growth in the absence of AZT. Data represent n=3 independent biological colony isolates plated in technical duplicates. Fractional survival is shown as mean and standard deviation. Representative images of plates are shown. Colony counts and all calculations are provided in the Supplemental Data file.

In the *ΔholC* background (Figure 9A), cells carrying the empty vector were highly sensitive to AZT, with survival reduced at 12.5 ng mL^-1^ and further reduced at 25 ng mL^-1^. Expression of χ alone resulted in the highest survival among the constructs at both AZT concentrations, although survival declined at the higher AZT concentration. In contrast, expression of ψ alone, the ψ-GS12-χ fusion, or the ψχ operon produced survival values similar to the vector control at both AZT concentrations tested, indicating no improvement in AZT tolerance.

In WT cells (Figure 9D), survival declined at both AZT concentrations tested. Survival values were similar among constructs at 12.5 ng mL^-1^ AZT. At the highest concentration tested (25 ng mL^-1^), ψ-GS12-χ showed the lowest survival relative to the vector control and other constructs. Expression of χ alone modestly improved survival at 12.5 ng mL^-1^ relative to the WT empty-vector control. Importantly, the observation that χ alone improved survival in both the Δ*holC* and WT backgrounds while ψ-containing complexes did not, indicates that AZT tolerance requires a pool of χ that functions outside the canonical clamp loader.

Total viable counts measured in the absence of AZT showed similar values for most constructs in both *ΔholC* and WT backgrounds (Figure 9B, E). In *ΔholC* cells, expression of the ψχ operon increased baseline viable counts relative to the empty-vector *ΔholC* strain, whereas the other constructs produced similar values. In WT cells, viable counts were similar across all constructs. Variability between replicates prevented the detection of statistically significant differences between conditions; accordingly, no statistically significant differences were observed among the constructs under any of the conditions tested.

Representative serial dilution plates (Figure 9C, F) reflected trends quantified in panels A through D. However, colonies expressing ψ-GS12-χ are smaller in both *ΔholC* and WT backgrounds, highlighted in the zoomed −4 and −5 dilutions in Supplemental Figure 6. Although this small colony phenotype was visually apparent, it did not reduce quantified viable-count measurements.

### Dominant-Negative Effects of ψ-GS12-χ Fusion Depend on χ-SSB Binding

The reduced colony size observed for ψ-GS12-χ expressing strains suggested that the fusion may interfere with normal cellular processes. Consistent with this interpretation, expression of ψ-GS12-χ in WT cells gave small colonies even in the presence of WT ψχ (Figure 9F and Supplemental Figure 6) and reduced survival at the highest AZT concentration tested (Figure 9D). Introduction of the χR128A mutation in ψ-GS12-χ, which disrupts χ-SSB binding, partially restored survival to AZT in WT cells, while expression of the ψχ operon produced intermediate behavior. Additionally, the small colony phenotype was reversed in both WT and Δ*holC* cells when the χR128A mutation was introduced into ψ-GS12-χ. Together, these findings demonstrate that the ψ-GS12-χ fusion had a dominant negative effect on cell growth in both Δ*holC* and WT backgrounds (Figure 9), with the strongest effects observed under AZT stress. Disruption of the χ-SSB interaction partially alleviated these defects, underscoring the importance of regulated χ-SSB binding for cell growth and DNA damage tolerance. While the expression of the fusion gives a different phenotype than expression of WT ψχ, neither is able to substitute for χ alone in AZT tolerance suggesting the fusion may still serve as a useful tool to probe which χ-containing complexes are important in different cellular contexts.

## DISCUSSION

The χ protein has a well-established role as a subunit of the DNA pol III HE, the main replicase in *E. coli*. Chi is an accessory subunit of the clamp loader portion of the holoenzyme and has no enzymatic activity on its own. Its primary role is to mediate physical interactions between the holoenzyme and SSB, which binds ssDNA that forms on the lagging strand during replication. Chi-SSB interactions stabilize the replicase on DNA, promote efficient DNA synthesis on SSB-bound DNA, and help remodel SSB on DNA to facilitate clamp loading (9–12). In addition, χ has been implicated in DNA damage tolerance to AZT, a drug that stalls replication forks (17). Although χ is required for AZT tolerance, its precise mechanistic role remains unclear. Notably, AZT tolerance also requires the YoaA helicase, which interacts with χ in a complex that is independent of DNA pol III HE. This raises the possibility that χ contributes to tolerance either through its role in the clamp loader, its interactions with YoaA, or potentially through associations with other unidentified partner proteins. Consequently, it remains unclear which χ-containing complex is most crucial for function in vivo.

To distinguish the function of χ as subunit of the DNA pol III HE from its role within the YoaA complex, we engineered a ψ-χ fusion designed to be uniquely incorporated into the clamp loader without altering the free pool of χ that can associate with the helicase. Because the χ binding surfaces for ψ and YoaA overlap (14, 17), point mutations cannot selectively disrupt one complex without affecting the other (14, 17). Consistent with this overlap, Y2H analysis confirmed that ψ and YoaA compete for binding to χ (Figure 7), supporting the use of the ψ-GS12-χ fusion strategy to restrict χ to clamp loader-associated complexes.

The design and function of amino acid linkers are common strategies in protein engineering to preserve protein structure, function, and interactions (41). Gly-rich linkers, in particular, are naturally occurring separators that connect domains within proteins while allowing them to retain discrete functions (31–34). These flexible linkers help stabilize interactions between protein partners, particularly when interactions are weak, transient, or involve disordered regions. Gly-Ser linkers are also relatively hydrophilic, which favours extended conformations over collapsed ones. By minimizing steric hindrance, they allow proper folding and function of the fused domains. In this study, our goal was to create a Gly-Ser linker between ψ and χ, providing sufficient flexibility to support stable domain interactions without compromising function both within the clamp loader or DNA pol III HE.

Both ψ-χ fusion proteins were expressed with an N-terminal His-tag and purified by chromatography. The His-tagged ψ-GS8-χ eluted from immobilized metal ion affinity chromatography in two separate peaks. SDS-PAGE analysis showed the two peaks contained the same polypeptide, suggesting that ψ-GS8-χ may adopt different conformations, folding states, or oligomeric states. Additionally, ψ-GS8-χ was poorly soluble and precipitated in several purification and buffer conditions, suggesting misfolding or aggregation. These observations indicate that the GS8 linker may be too short to maintain the correct intramolecular orientation between ψ and χ. As a result, the fusion may fail to form a stable heterodimer structure, as predicted by AlphaFold, and hinder the high-affinity binding of ψ to χ (Figure 10). When ψ is overexpressed alone, the protein is insoluble, suggesting that loss of ψ-χ contacts could further reduce solubility (40). Alternatively, oligomers may form if ψ from one fusion protein binds χ from another, generating higher-order assemblies that are less soluble (Figure 10).

**Figure 10:**
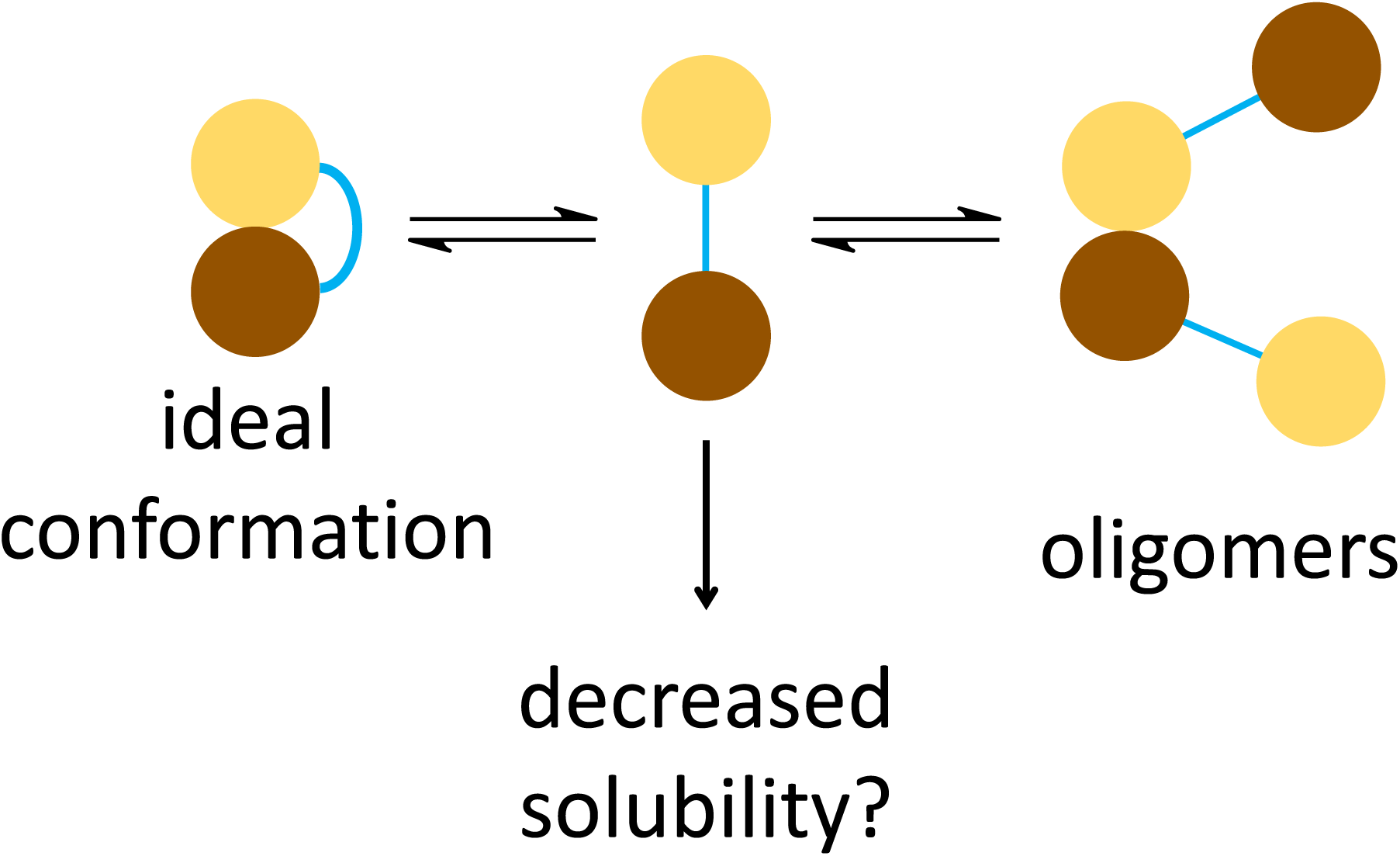
Model of fusion stability. Schematic representation of proposed outcomes for ψ-χ fusions. In the ideal conformation, ψ and χ interact intramolecularly to form a stable complex resembling the native heterodimer. If the fusion prevents proper ψ-χ association, the protein may adopt a misfolded or partially unfolded state with decreased solubility. Alternatively, ψ and χ domains from different fusion molecules may interact intermolecularly, generating oligomeric assemblies that further reduce solubility.

In contrast, the ψ-GS12-χ fusion protein eluted as a single peak from the immobilized metal ion affinity chromatography and was considerably more soluble than ψ-GS8-χ. Further purification by cation exchange yielded a doublet of closely spaced peaks, which may reflect conformational heterogeneity arising from the disordered region of ψ. CD spectroscopy confirmed that ψ-GS12-χ retained secondary structure similar to WT ψχ (Figure 4C), although CD cannot rule out multiple conformational states as illustrated in Figure 10.

Importantly, DSF showed that both SP fractions of ψ-GS12-χ had identical T_m_ values, indicating that any conformational variability does not compromise stability. Moreover, the overall T_m_ of ψ-GS12-χ was significantly higher than that of WT ψχ, indicating a clear stabilizing effect of the GS12 linker (Figure 4A-B). This stabilization likely arises from improved spatial accommodation between the two domains, with the flexible linker relieving strain and allowing native-like intramolecular interactions when compared to the GS8 linker. The fusion also ensures that ψ and χ are always held close in space, so when they transiently dissociate, they can rebind more rapidly through intramolecular association, which may further enhance stability compared to WT ψχ.

DSF analysis also showed that the Peak 1 of ψ-GS8-χ had tight error bars for T_m_, indicating a reproducibly stable population, whereas Peak 2 displayed larger variability (Figure 4A). This suggests that a fraction of the ψ-GS8-χ population adopts a reproducibly stable conformation. However, the broader variability and reduced solubility of the GS8 fusion overall indicate that the shorter linker may not reliably support the correct folding or prevent aggregation.

Despite these structural differences, both fusion proteins retained functionality in the context of the clamp loader. ATP hydrolysis is a critical step in the clamp-opening cycle, driving the conformational changes required to open and release the beta clamp from DNA. In our assays, observed ATP hydrolysis rates for clamp loaders containing ψ-GS8-χ or ψ-GS12-χ were similar to those of WT (Supplemental Figure 3), in the presence and absence of SSB, indicating that incorporation of the fusions does not impair ATPase activity to the extent that it becomes rate-limiting under steady-state conditions. However, for the WT clamp loader, ATP hydrolysis is not the rate-limiting step in steady-state turnover. The observed rate is instead limited by a downstream step in the clamp-loading pathways that is not affected by SSB, as also observed for the χ-deficient clamp loader. Consequently, steady-state ATPase measurements do not provide a sensitive readout of χ-SSB interactions or their potential disruption in the fusion constructs but instead report on the integrity of the core ATP-driven motor.

To examine clamp loading more directly, we measured clamp-closing kinetics using stopped-flow under single-turnover conditions (Figure 5A-C). These experiments isolate the clamp-closing step and are sensitive to χ-SSB interactions. Although the rate of clamp-closing was similar to WT, the fluorescence amplitude was reduced for both fusions, particularly ψ-GS8-χ (Figure 5A-C). The *K_d_* values for clamp binding (Figure 6A-C) and opening kinetics (Supplemental Figure 4) were unchanged, suggesting that the reduced amplitude is not due to differences in clamp binding or opening. Moreover, equilibrium binding experiments showed that the fraction of clamp loader-clamp complexes in the open conformation was similar to WT. These observations suggest that the reduced amplitude instead reflects subtle changes in conformational dynamics under the specific assay conditions.

Overall, these findings confirm that the ψ-GS12-χ fusion is structurally stable, soluble, and functionally competent within the clamp loader. However, clamp-loading and ATP hydrolysis assays test only core biochemical activities and do not reveal whether the fusion can support specialized roles at the replication fork, such as those required under DNA-damage stress. To address this gap, we next examined whether ψ-GS12-χ could substitute for χ in vivo, focusing on its role in AZT tolerance.

The expression of either the ψ-GS12-χ fusion or the WT ψχ operon did not rescue the AZT sensitivity of Δ*holC* cells (Figure 9A). In contrast, expression of χ alone restored survival beyond WT levels across AZT concentrations, although survival still declined as drug concentration increased (Figure 9A and D). This demonstrates that the χ associated with the replicase is unlikely contributing to AZT tolerance. Instead, χ alone is needed to support AZT tolerance. This may reflect the need for the pool of χ that interacts with YoaA, or possibly with other unidentified binding partners, rather than clamp loader-associated χ. Importantly, the inability of ψ-GS12-χ to restore AZT tolerance does not mean that the fusion is nonfunctional in vivo, as immunoblot analysis confirmed expression of the fusion under the induction conditions used (Figure 8). Rather, these results suggests that different χ-containing complexes may serve distinct roles. The key question is therefore not only which χ-containing complex is most important, but also where χ exerts its essential functions in the cell.

In WT cells, expression of ψ-GS12-χ produced fractional survival similar to that of the WT operon under AZT conditions (Figure 9D), with both showing reduced survival relative to the vector control. However, these reductions in fractional survival were not statistically significant. This suggests that the reduced fractional survival arises from the overexpression of ψχ-containing complexes rather than from a fusion-specific effect. Notably, expression of the ψ-GS12-χ produced a small colony phenotype (Figure 9C and 9F). Introduction of the χR128A mutation, which disrupts the χ-SSB interface, reduced the small colony phenotype in both WT and *ΔholC* cells (Figure 9F and 9C). These findings support the idea that χ-SSB binding is essential for maintaining normal cellular function.

Taken together, these findings indicate that while ψ-χ fusions are structurally stable and biochemically competent in vitro, their overexpression in vivo can be detrimental to cellular growth, as evidenced by the small colony phenotype observed in both WT and *ΔholC* backgrounds (Figure 9 and Supplemental Figure 6). This phenotype may arise, in part, from incorporation of the fusion into the clamp loader, where altered ψ-χ connectivity could subtly impair replication dynamics. This phenotype may also be linked to the stability and assembly properties of the fusion. Although ψ-GS12-χ was more stable than ψ-GS8-χ, the enforced tethering of the two subunits may still favor non-native assemblies. The fusions may also assemble into clamp loader subcomplexes that are not part of the full DNA pol III HE, consistent with evidence that clamp loader components can exist independently (9, 42, 43). These subassemblies, or free ψ-GS12-χ protein, could engage SSB or DNA in ways that compete with canonical replisomes (44, 45).

Because only a limited number of DNA pol III complexes exist in each cell, not all ψ-GS12-χ fusions are expected to be incorporated into holoenzymes (46). A substantial fraction of the fusion protein is therefore likely free, where it may bind SSB inappropriately. Such improper binding could increase its local concentration at replication forks and interfere with normal protein handoffs. A similar effect may also occur upon overexpression of the WT ψχ operon, suggesting that elevated levels of ψχ complexes, rather than the fusion alone, can perturb normal SSB-mediated protein exchange. However, by tethering ψ to χ, the modularity of the clamp loader is reduced, which could further exacerbate these effects by trapping χ on SSB and preventing access for other SSB-interacting proteins (SIPs) such as RecO, RecQ, PriA, or YoaA (47–50). In WT cells, χ normally engages SSB dynamically, allowing orderly exchange with downstream repair and restart factors. In contrast, the ψ-χ fusion could interfere with SIP recruitment, as supported by the dominant-negative phenotypes in vivo and the partial rescue by the χR128A mutation, which weakens χ-SSB binding. Such dominant-negative effects are consistent with established models in which overexpression of SSB-interacting proteins sequesters SSB and disrupts replication-associated protein exchange (44, 45).

Additional mechanisms may also contribute. Elevated ψχ levels may alter clamp loader stoichiometry, titrate essential factors away from the replisome, or promote higher-order assemblies if the disordered region of ψ is engaging improperly. Persistent sequestration of clamp loader subunits, or free ψ-GS12-χ on SSB-DNA, could further reduce their availability elsewhere in the genome, slowing replication and increasing transcription-replication conflicts.

This model is further supported by live-cell imaging studies showing that SSB foci colocalize with DnaQ, DnaE, HolC, and HolD at replication forks (>90% colocalization), consistent with their established role as part of the gamma complex (51). These data reinforce that ψ and χ function directly at active replisomes where SSB is concentrated, providing the spatial context for our model. The dominant-negative effects of the ψ-GS12-χ fusion are therefore most likely to occur at replication forks, where persistent χ-SSB binding would block dynamic handoffs to other SIPs, including YoaA. Failure to recruit YoaA explains why neither the ψ-GS12-χ fusion nor the WT operon restored AZT tolerance in *ΔholC* cells, whereas expression of χ alone was sufficient.

The importance of this dynamic exchange is especially evident under AZT stress, which generates ssDNA gaps behind replication forks. Chi has been implicated in recruiting the YoaA helicase, which may facilitate gap processing and enhance AZT tolerance. Consistent with this, deletion of *yoaA* improves growth and viability in *holC* mutants, indicating that YoaA activity is toxic when χ is absent (17). These genetic data suggest that χ may be essential for regulating or stabilizing YoaA, and that its absence leads to deleterious outcomes. The fusion may disrupt this regulatory role by anchoring χ to ψ and SSB, limiting its ability to engage YoaA when needed. Importantly, both the fusions and the WT operon are not restricted to clamp loader incorporation and may also exist as independent pools in the cell. In this context, persistent SSB binding by free WT or fusion complexes could further reduce opportunities for YoaA-χ interactions, providing a mechanistic explanation of why neither the ψ-GS12-χ fusion nor the WT operon restored AZT tolerance in Δ*holC* cells, whereas expression of χ alone was sufficient. In contrast, the R128A mutation reduces the χ-SSB occupancy, allowing YoaA or other SIPs to access SSB and partially restore tolerance.

Furthermore, the sequestration of clamp loaders on SSB-DNA may also diminish the availability of functional complexes elsewhere in the genome, slowing replication and increasing transcription-replication conflicts. These broad cellular effects are consistent with the impaired growth phenotypes observed even without added stress.

We propose a model in which modular interactions between χ, ψ, and SSB must remain flexible and tightly regulated to support replication. When this regulation is disrupted - whether by fusing χ to ψ, by altering expression levels, or by other perturbations - the modularity is lost, leading to inappropriate or persistent SSB engagement that disrupts normal clamp loader function and impairs DNA damage tolerance. Disruption of the χ-SSB binding interface through the χR128A mutation in the fusion alleviated these defects, highlighting the importance of regulated χ-SSB dynamics in preserving clamp loader integrity and replication fork stability.

## DATA AVAILABILITY

Expression plasmids for proteins used in this study will be shared upon request to the corresponding author.

## SUPPLEMENTARY DATA

Supplementary Data are available at NAR online.

## AUTHOR CONTRIBUTIONS

K.A.P.P., S.T.L., and L.B.B. conceived and designed the project. K.A.P.P., J.D.G., M.J.P., and E.S.P.N. collected the data. K.A.P.P and L.B.B performed the analysis. K.A.P.P wrote the original draft. K.A.P.P, J.D.G., M.J.P., E.S.P.N., S.T.L., and L.B.B. reviewed and edited the draft. S.T.L. and L.B.B. supervised the project. L.B.B acquired funding for the project.

