## Supplemental Figure for "A Novel ψ-χ Fusion Protein for Unravelling the Contributions of χ to DNA Replication and Repair"

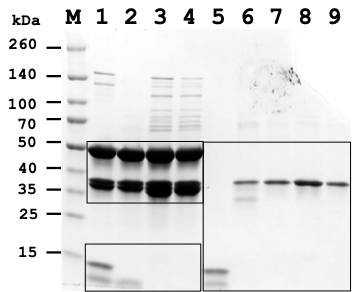

**Supplemental Figure 1:** SDS-PAGE analysis of purified ψ-χ fusion proteins and γ complex variants. A) Coomassie-stained 12% SDS–PAGE gel showing purified clamp loader complexes and ψχ fusion proteins. Lane M, molecular weight marker. Lane 1, WT γ complex; lane 2, γ complex Δχ; lane 3, γ complex ψ-GS12-χ; lane 4, γ complex ψ-GS8-χ; lane 5, ψ-GS8-χ #2; lane 6, ψχ; lane 7, ψ-GS12-χ #1; lane 8, ψ-GS12-χ #2; lane 9, ψ-GS8-χ #1. Boxes indicate the sections of the gels used in Figure 3.

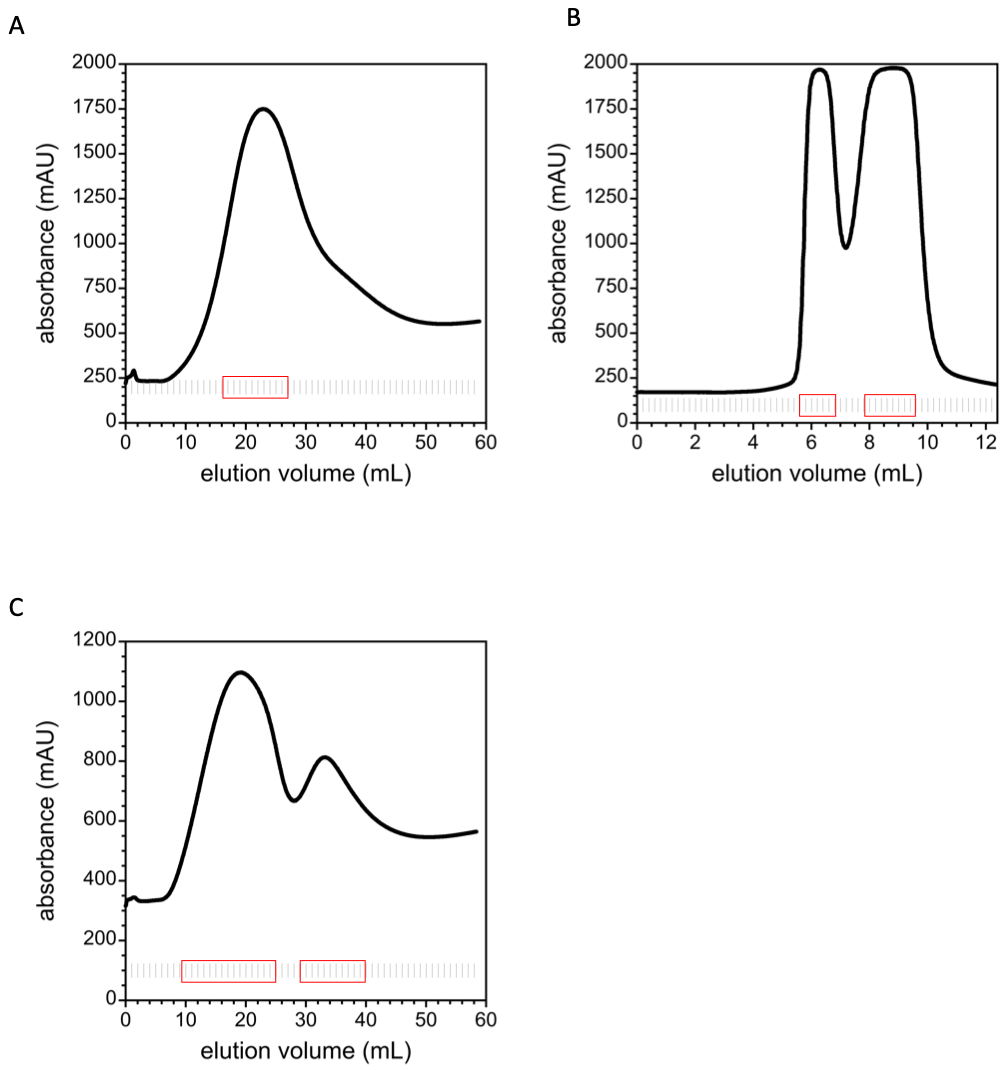

**Supplemental Figure 2:** Purification of fusion proteins. UV absorbance profiles for ψ-GS12-χ from HisTrap FF (A) and SP HP (B) columns are shown, with collected fractions indicated by vertical lines and pooled fractions by red boxes. The UV absorbance profile for ψ-GS8-χ from the HisTrap FF column is shown in (C), with fractions indicated as above. All chromatograms are representative.

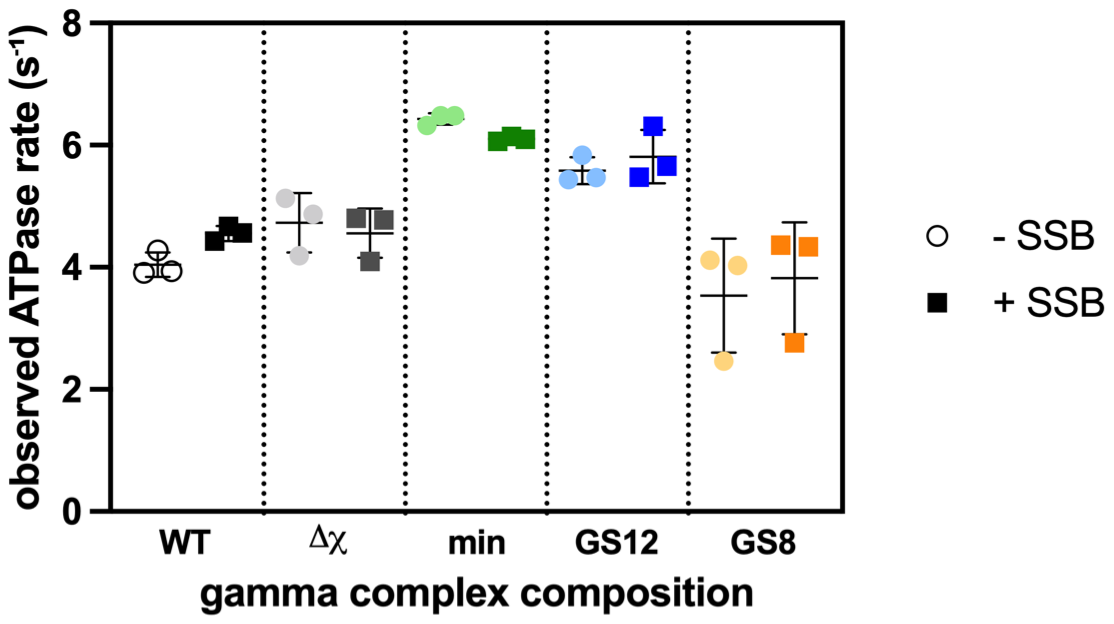

**Supplemental Figure 3:** Steady-state ATP hydrolysis by γ complex variants. ATPase activity was measured using an enzyme-coupled assay monitoring NADH oxidation at 340 nm. Reactions contained 0.5 μM 20-bp duplex DNA with a 5’ dT65 overhang, 80 nM γ complex, 1 μM β clamp, 0.5 mM ATP, and were performed in the presence (squares) or absence (circles) of 0.75 μM SSB as indicated. ATP hydrolysis rates were determined for WT γ complex, γ complex lacking χ (Δχ), a minimal complex lacking χ and ψ (min), and γ complexes containing the ψ-GS12-χ or ψ-GS8-χ fusion proteins. Rates were calculated from the decrease in NADH absorbance and normalized to γ complex concentration. Points represent individual measurements and bars indicate mean ± SD.

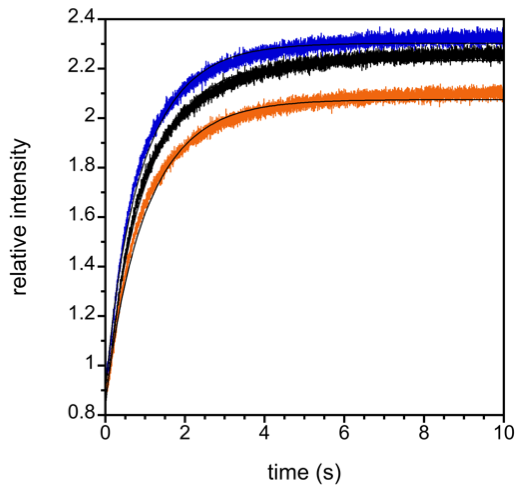

**Supplemental Figure 4:** Clamp opening by γ complex variants containing fusion proteins. Clamp opening reactions containing 60 nM γ complex clamp loader, 40 nM β-TMR_2_, and 0.5 mM ATPγS. Gamma complex variations are shown as WT (black), ψ-GS12-χ (blue), and ψ-GS8-χ (orange). Solid black lines through the data show fits to a double exponential (Equation 5). n = 1

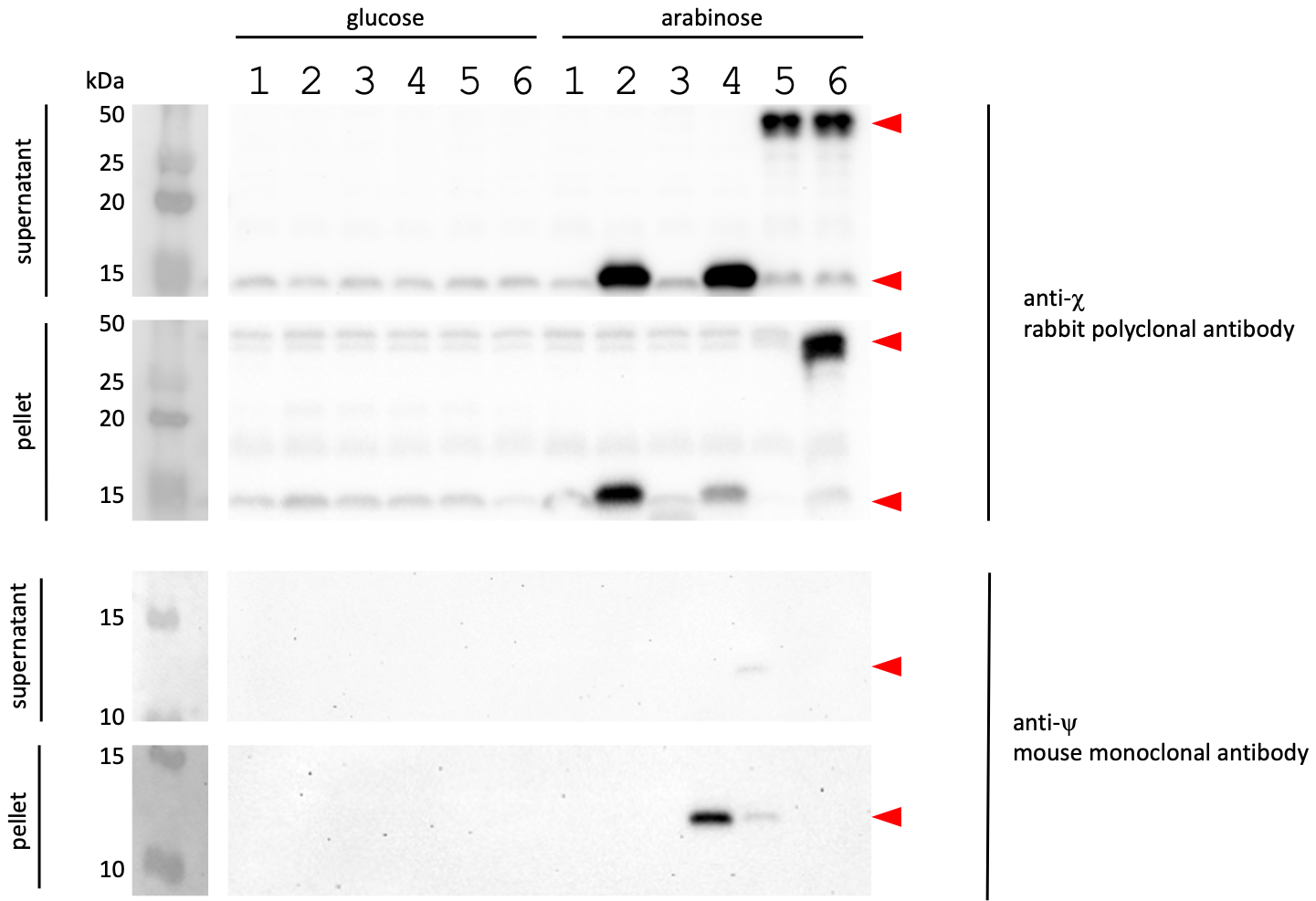

**Supplemental Figure 5. Uncropped Western blots corresponding to Figure 8.**
Uncropped immunoblots showing expression of χ, ψ, and the ψ-GS12-χ fusion constructs under complementation conditions. WT *E. coli* cells were grown under the same induction conditions used for complementation assays and fractionated into soluble (supernatant) and insoluble (pellet) fractions prior to analysis. Equal amounts of total protein were loaded per lane based on A280 quantification. Blots were probed with anti-χ rabbit polyclonal or anti-ψ mouse monoclonal antibodies, as indicated. Lanes 1-6 were grown in glucose (repressed conditions) and lanes 7-12 in arabinose (induced conditions). Molecular weight markers are shown on the left, and arrowheads indicate the positions of χ, ψ, and ψ-GS12-χ.

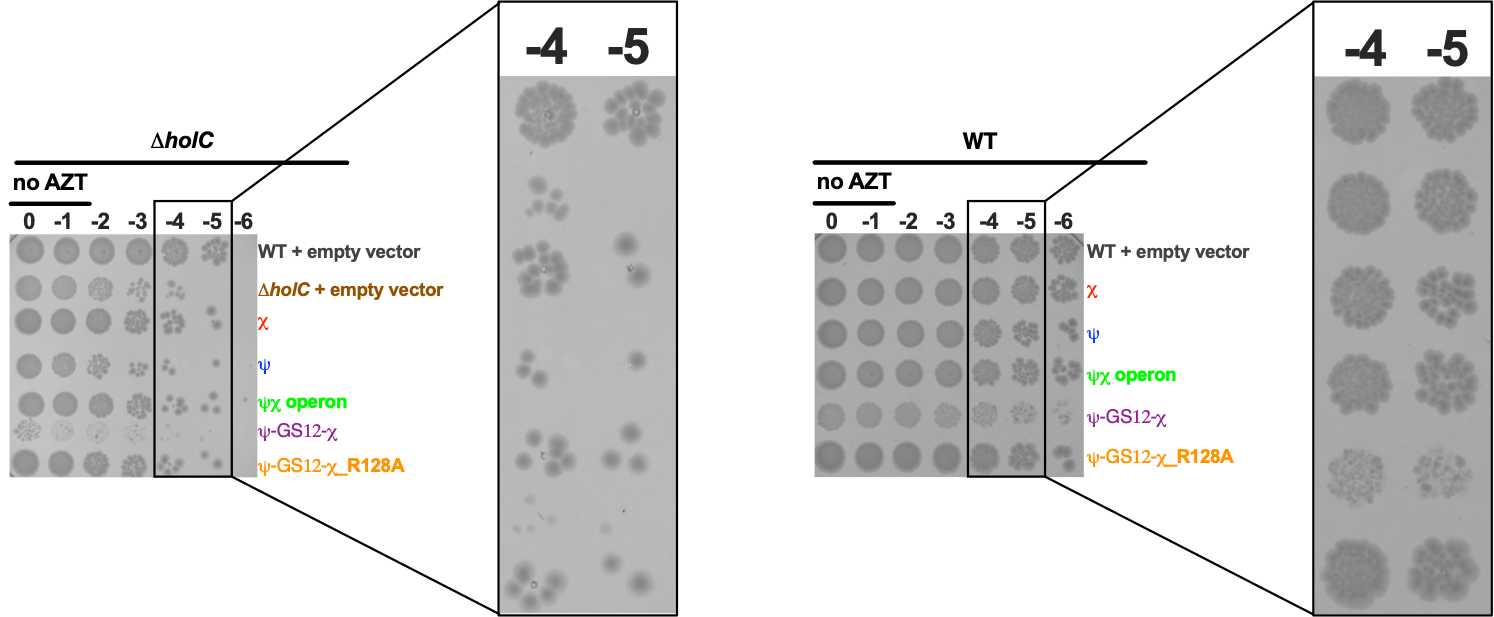

**Supplemental Figure 6. Zoomed view of serial dilution spot assays highlighting colony size phenotype in Figure 9.** Representative serial dilution spot assays from Figure 9 showing WT and Δ*holC* cells in the absence of AZT. Boxes indicate the −4 and −5 dilutions, which are shown at higher magnification to better visualize colony morphology. Images correspond to those shown in Figure 9.
